# Network topology and evolution of the gene co-expression of T-cells during immuno-senescence

**DOI:** 10.1101/2021.09.01.458500

**Authors:** Megan L. Mair, Nicolas Tchitcheck, Tarynn M. Witten, Véronique Thomas-Vaslin

**Affiliations:** Virginia Commonwealth University, Center for the Study of Biological Complexity; Department of Computer Science US-VA 23284-2030 Richmond, United States; Sorbonne Université, INSERM, Immunology, Immunopathology, Immunotherapies, UMRS959, F-75013 Paris, France

## Abstract

To better understand the potential impact of the gene expression network structure on the dynamics of immune-senescence and defects of cell functions during aging, we investigated network structures in both young and old individuals. We analyzed the gene co-expression networks (GCNs) derived from an aging signature of 130 immune-related genes obtained from CD3+ T-cell splenocytes extracted from FVB/N, C57BL/6N, and BALB/c mice at ages 2 and 22- 24 months. The network structure for the two different mouse age-groups was derived and subsequently analyzed. Analysis of network hubs using clustering coefficients, degree, betweenness, eigenvector, and closeness centralities, as well as local, indirect, and total influence measures, demonstrated changes in gene behavior and network control between the two age groups. Our quantification shows that the young, 2-month old mouse network is more organized than the 22-24-month, old mouse network, while the network structure of the older mouse GCN appears to be far more complicated but far more dispersed. Changes in network structure between the old and young mice suggest deterioration in transcription regulation with age in peripheral T-cells, particularly within the TCR signaling pathway, and potential compensatory mechanisms in older T-cells to overcome loss to regular function resulting from transcriptional irregularity. These results demonstrate the need for more research into gene co-expression in peripheral T-cells in order to better understand both network irregularities and the phenotypic dysfunction observed in older individuals.

**Author Summary:** In order to better understand the potential mechanisms of transcriptional irregularities in the immune system with aging, we analyzed the structure of gene co-expression networks of T-cells extracted from the spleens of 2 and 22-24-month old mice. Gene co-expression describes the correlation relationship between two expressed genes; as the expression of one gene goes up, the expression of another gene might also increase (or, conversely, decrease). Strong gene co-expression relationships can signal the existence of a number of important biological phenomena, such as two genes belonging to a transcription pathway or protein structure. Network diagrams visualizing these co-expression relationships in both younger and older mice demonstrated the existence of differences in network structure and properties that may be attributed to the aging of the immune system. Network mathematical methods were used to examine the complexity of each network. We found that the younger mouse network was more organized than the older mouse network. The older mouse group exhibited a 255% increase in co-expression relationships but a decrease of 92% of the connections from the young mouse network. This suggests the older mouse T-cells suffer dysfunction at a transcriptional level. This results in the loss of regular immune and cellular functions. These results demonstrate the importance of future research into gene co-expression to decipher senescence or diseases that perturb gene expression through time.

## Introduction

Aging is a natural process that progressively alters biological cell functions at the microscopic level with cell senescence, and extends up to the organ level and overall function of the organism, leading to complex disorganization at various scales in living systems [1]. The network connectivity between the nodes composing the multi-level hierarchical network of “aging” could thus be altered at various biological levels. This could lead to rupture of stability and robustness of dynamic interweaved networks with age.

At the mesoscopic level, the population of individuals aged 65 and over is increasing [2]. The economic and biomedical influence of the elderly population will have a significant impact upon the global economy [3–5]. Chronic inflammation and susceptibility to infection are traits of immuno-senescence, the term used to describe the overall age-related changes of the immune system [6–10]. Immune dysfunction in the elderly contributes to increased susceptibility to infection and inflammation [11–13]. Immuno-senescence can have a negative impact on health outcomes including morbidity and mortality in older adults [6, 10–18]. This was recently underlined with the Covid-19 pandemic, where older individual were more susceptible to severe disease and mortality [19].

Aging is often characterized by perturbations and/or remodeling of the T-lymphocyte system [20–23]. Aging contributes to the decrease of naïve T-cell production [24], first by thymic involution starting at puberty. These perturbations of production are then compensated by increasing the selection and the proliferation of effector/memory T lymphocytes populations, which were previously exposed to an antigen. As the organism ages, the accumulated effector/memory cells often become senescent non–functional T-cells. These fill the “immunological space,” replacing the decreasing population diversity of naïve cells, which are able to recognize and combat sources of new antigens.

Immuno-senescence also affects mice; mice are thus a good model to study the dynamics of T lymphocyte aging through micro (cells) and macro (tissues/organs) levels. Mice have previously been used to demonstrate other changes in T lymphocyte biology in the context of aging. For example, partial clonal deletion is known to occur in the spleen in old mice. Previously,

Thomas-Vaslin *et al*. quantified the heterogeneity of T-cell population proportions based on their proliferation rates, according to various mice genetic origins [25]. This revealed decreased proliferative and renewal capacities of T-cells with aging. These selection processes also result in a diminished T-cell receptor (TCR) repertoire biodiversity, required for the recognition of new antigens [23]. Thomas-Vaslin *et al*. have additionally demonstrated that T-cell homeostasis is maintained in young mice after a transient perturbation [26], with a biodiversified repertoire. However, at mid-life the immuno-senescence decreases the turnover and proliferation of naive T-cells with new TCRs, while memory T-cells and oligoclonal expansion accumulates. The consequences of the natural immunodepression during aging could be exacerbated by the effects of transient perturbations such as immunosuppressive treatments which kill dividing cells, like chemotherapy [25]. Most immunopathologies in humans also drive oligoclonal expansions of T-cells [27].

These observations suggest that the T-cell dynamic network interactions and regulation processes could be influenced by age. Our hypothesis is that responses to molecular perturbations, from the level of gene transcription, is optimized in the T-cells of young mice, and that this robust response is diminished in old mice.

Understanding the complexity of gene expression and cell pathways can be approached through the analysis of network gene co-expression relationships [28]. Gene co-expression networks have been used to study the biology of organisms such as plants, mice, and humans [29–31]. Gene co-expression describes simple correlation relationships between the expression levels of two genes. For example, as the expression of one gene increases, a given gene might have a strong positive or negative correlation with the expression of a second gene. Strong gene co-expression relationships correlate with a number of intracellular biological processes, including transcription pathways, protein complex formation and cell signaling. Changes in co-expression relationships and subsequent network topology can therefore provide insight into altered biological pathways, for instance, in the contexts of disease or aging [28, 32–33]. Gene co-expression analysis through network structures can begin to provide a more holistic view of biological processes related to gene expression, allowing for the detection of changes not otherwise evident in simple gene expression data.

In order to understand these more complicated interactions, mathematical, bioinformatics and computational methods may be brought to bear. Among the system-level tools in these fields are those of network analysis. There are already a number of historical research papers that have applied network analysis techniques to studying various aging processes and variation of connectivity during aging of various species [34–45], up to the theory of aging networks. Network comparison techniques allow for the detection of potential differences in gene co-expression across a variety of biological conditions, including possible age-related changes in gene co-expression.

In this paper, we present an analysis of aging-related, dynamical transcription modification using gene co-expression data derived from peripheral T-cells from both young (2 months old) and old mice (22/24-months old). These gene co-expression networks were derived from a shared signature of 130 immune-related genes obtained from CD3+ splenocytes extracted from mice strains that display genetic variability (FVB/N, C57BL/6N, and BALB/c) [46]. We applied a number of network analysis methods to determine the existence of optimal network capacities and connectivity in young mice. We then investigated the age-related changes in network properties and structures by analyzing the loss of gene co-expression and the newly gained gene co-expression relationships in the older mouse network. We discuss how these changes may be associated with mouse species immuno-senescence, and how these changes might give insights into the aging of the human immune system.

## Results

### General terminology of structure of gene co-expression networks

We investigate here the topology and evolution of gene co-expression networks (GCN), across young and old mice, tracking the lost and gained gene co-expressions and the resulting remodeling of the structure of the networks that could influence the possible robustness/efficacy/resilience of the adaptive immune cells. For the purposes of our upcoming discussion, a node/vertex/hub means a gene that is expressed as an mRNA and an edge means that there is a single link, undirected, between two expressed genes in the network (*i.e*., there exists a strong correlation between the expression of the two linked genes). A reference for important network terminology can be found in Table 1, and important concepts are illustrated in Fig 1.

**Figure 1.**
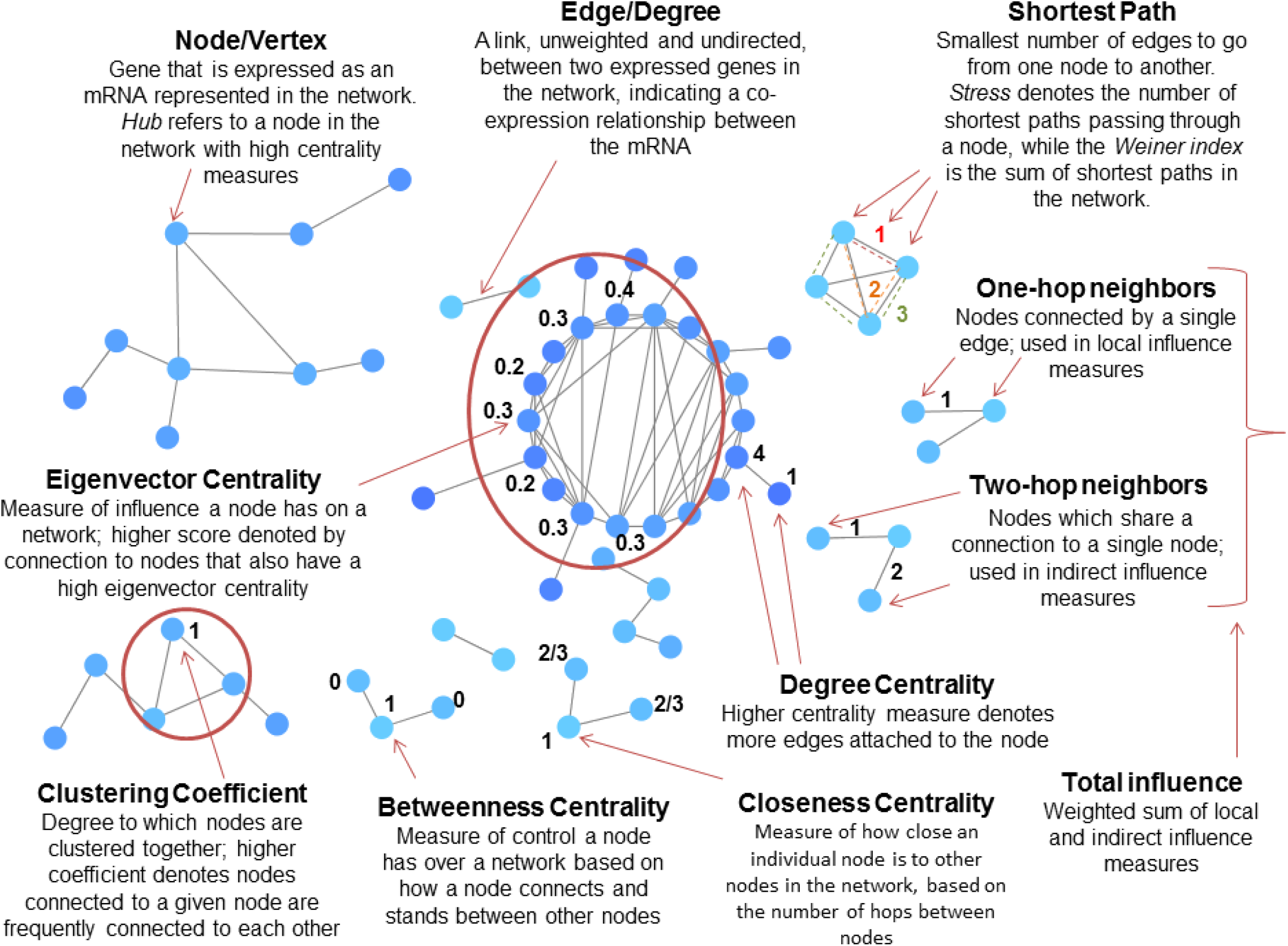
Diagram illustrative of key network terminology. Terminology used in this paper is important for the discussion of Gene Co-expression Networks (GCNs).

**Table 1.** Table of network terminology.

| **Term** | **Definition for gene co-expression network (GCN)** |
| --- | --- |
| **Node/Vertex/Hub** | Gene that is expressed as an mRNA represented in the network |
| **Edge** | A link, undirected, between two expressed genes in the network, indicating a co-expression relationship between the mRNA |
| **Degree centrality** | Centrality measure denoted by the number of edges directly attached to a node |
| **Eigenvector centrality** | Measure of influence of a given node in a network. A higher eigenvector centrality score means a node is connected to many other nodes with eigenvector centrality scores |
| **Closeness centrality** | Measure of how close an individual node is to other nodes in the network |
| **Betweenness centrality** | Measure of control a node has over a network based on how a node connects and stands between other nodes |
| **Eccentricity** | The maximum distance between a node and all other nodes |
| **Stress** | The number of shortest paths passing through a given node, indicates the importance of a node holding together a network |
| **Weiner index** | Sum of all the shortest paths in the network |
| **Local influence** | The influence a node has over its directly connected, one-hop neighbors |
| **Indirect influence** | The influence a node has on its two-hop neighbors |
| **Total influence** | Weighted sum of direct and indirect influence |
| **Clustering coefficient** | Measure of degree to which nodes cluster around a given node in the network |

### Aging signature and hierarchical clustering of mice according to gene expression in splenic T-cells

To decipher gene expression modifications across aging, we performed a transcriptomic profiling of CD3^+^ purified splenocytes from 12 young mice age 2 months and from 11 old mice ages 22-24 months. The ICA/GSEA method was used to identify a signature of 130 genes able to distinguish young from old mice. Interestingly, the age-related gene signature allows us to segregate the 3 mice strains, considering only the young mice. However, this signature cannot separate the mice according to the genetic origin of old individuals due to the high variation of gene expression (S1 Figure). This suggests a precise synchronization and organization of gene expression in development that could vary according to genetic origins, allowing for the optimal functionality of T-cells in young mice. However, a disorganization of gene expression appears in old mice that could no longer maintain their individual genetic signature. This suggests an introduction of stochastic events in gene expression. To verify this hypothesis, we then proposed topological analysis of the gene-co-expression network.

### General Network Structure of T-cell expression in the spleen of young and old mice

To further understand the aging alterations that could impact T-cell functionality based on our gene signature, we generated two gene co-expression networks (GCN) based on the young and old aged mice. The network of young mice represent the optimal activity and functionality of the pool of all splenic T-cells, based on a consensus of expression data. The network of old mice represent the resilient T-cells that continue to live in the old mice, while the natural immunodepression has already removed more than 80% of naïve T-cells while favoring the proliferation and accumulation of effector/memory T-cells in the spleen [24–25]. Based on these two networks, we performed topological analyses to compare their structures and functionalities. We used various centrality network measurements applied to each GCN.

We first analyzed the structure of the gene co-expression network in young mice and compared it to the network structure of GCN belonging to old mice. Genes expression was detected with probes that cover messenger RNA from different genetic regions, including multiple probes that detect particular TCR proteins encoded from stochastic gene rearrangements of VDJ genes. The mRNA represented by each node is expected to give rise to the functional protein interactions related to known intracellular pathways expressed in the sampled T-cells. S1 Table lists all of the gene transcripts found in both of the networks, along with known human orthologs and known gene functions. Network files featuring the genes that appear in each network, as well as their corresponding co-expression relationships, can be found in S2 Table (young mouse GCN) and S3 Table (old mouse GCN).

### Pathway Analysis: KEGG /REACTOME

Genes appearing in the young and old mouse network were enriched in several immune system-related pathways; of the network genes contained in MSigDB, 33% in the young mouse network and 21% in the old mouse network were enriched for the REACTOME Immune System pathway. Despite having fewer nodes total, a higher percentage of genes in the young mouse network were enriched in KEGG and REACTOME immune system-related pathways compared to the older mouse network. Pathways unique to the young mouse GCN were genes involved in the translocation of ZAP-70 and the phosphorylization of CD3 and TCR zeta chains, which was significant at the 0.05 level. The old mouse GCN, on the other hand, had genes enriched for aldosterone-regulated sodium reabsorption and primary immunodeficiency, while the young mouse network did not. A complete summary of pathway enrichment results can be found in Fig 2.

**Fig 2.**
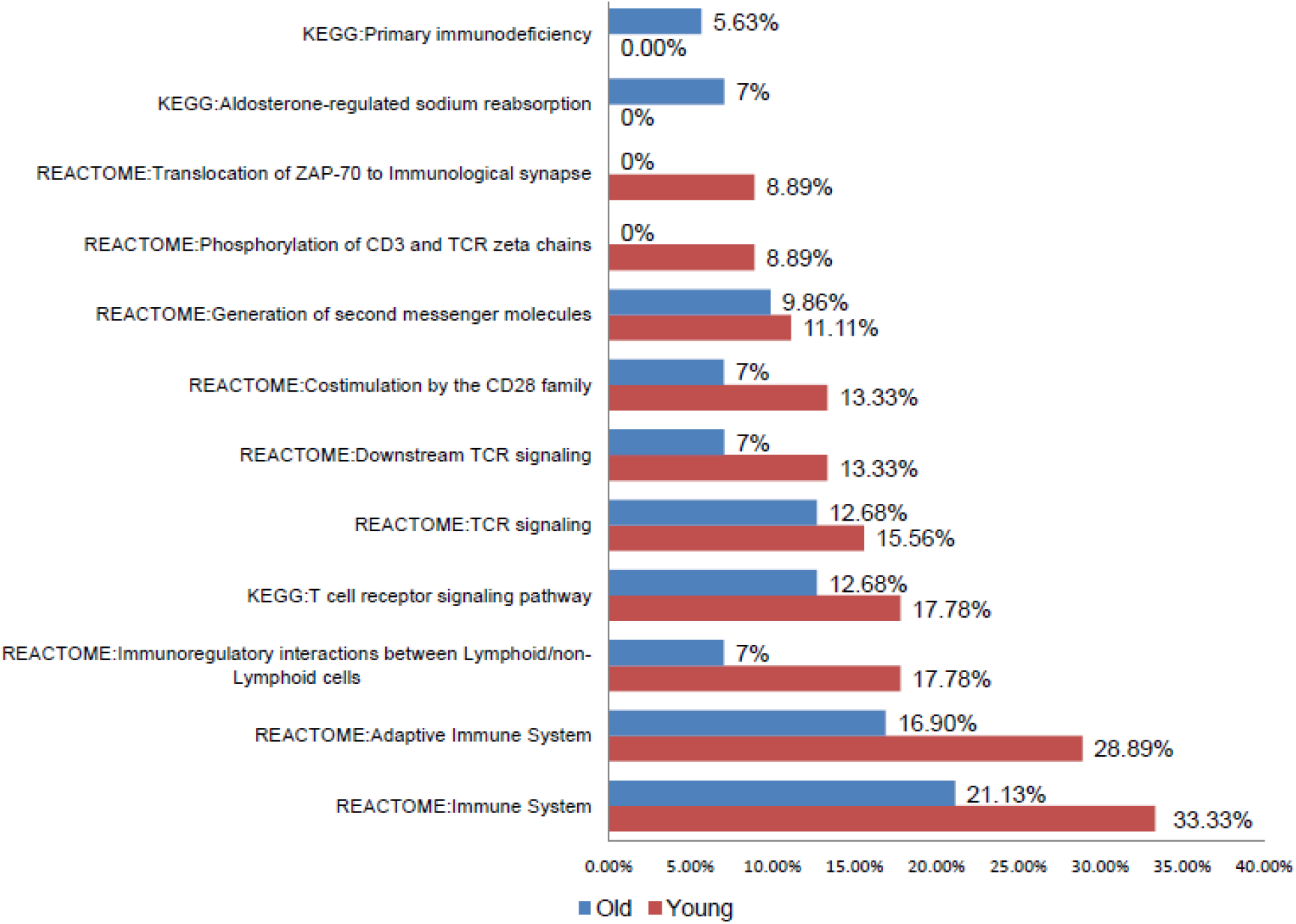
KEGG and REACTOME Pathway Enrichment. Percentage of genes enriched in each pathway in the young (red) and old (blue) mouse GCNs. Only the difference between the proportion of genes in the old and young networks enriched for involvement in the translocation of ZAP-70 and the phosphorylation of CD3 and TCR zeta chains was significant at the 0.05 level.

### Structure of the network in young and old mice as number of nodes and edges

The range of edges per node for the younger mouse GCN was 1-10 edges, with a mean edge count of 2.656. For the older mouse GCN, the range of edges per node was 1-15 edges per node with a mean degree count of 5.922 edges per node. For the 2-month old mouse GCN, the average node degree count falls within the range typical of a biological network [47]. This is not true for the 22/24-month old mouse network. A summary of some of these network differences can be found in Table 2. A t-test demonstrated a statistically significant difference between the mean edge counts of the two age-groups (*p <* 0.001).

**Table 2:** Summary of basic network statistics for the 2 and 22-24 month-old networks.

| Gene centrality measure | 2-month<br>GCN | 22-24-month<br>GCN | Difference |
| --- | --- | --- | --- |
| Mean Edge Count | 2.66 | 5.92 | Significantly different<br>$p < 0.001$ |
| Mean Shortest Path | 2.05 | 3.39 | Significantly different<br>$p < 0.001$ |
| Mean Clustering Coefficient | 0.25 | 0.34 | Significantly different<br>$p < 0.05$ |
| Weiner Index | 2206 | 76617 | N/A |
| Network Diameter | 7 | 12 | N/A |
| Core Network Radius | 4 | 6 | N/A |

Network attributes were highly altered across the two age-related networks. The 2-month GCN had a total of 64 nodes with 78 unique edges, while the 22/24-month old GCN had 102 nodes with 295 unique edges. A total of 53 nodes were found to be identical in both the 2-month and 22/24-month old networks. What is interesting is that despite this conservation of 53 nodes, only 7 edges are conserved across both networks. Table 3 lists the genes at the ends of each of the 7 edges. We highlighted these conserved edges in Fig 3. As we will later describe, the 7 highlighted conserved pairs in Fig 3 are ranked first or second in nearly all of the network centrality measures. Notably, most of the genes co-expressed and conserved between T-cells from young and old mice concern the TCR, which is required to receive signals from antigens and transduce signaling pathways to trigger various T-cell functions. However, the co-stimulation of the TCR involving CD28 and its downstream signaling [48] is disconnected from the core network gene expression in old mice, suggesting default of co-stimulation as in hallmark of senescent CD8 T-cells [49]. The overall network structure for both age-groups is summarized in an adjacency matrix in Fig 4.

**Fig 3.**
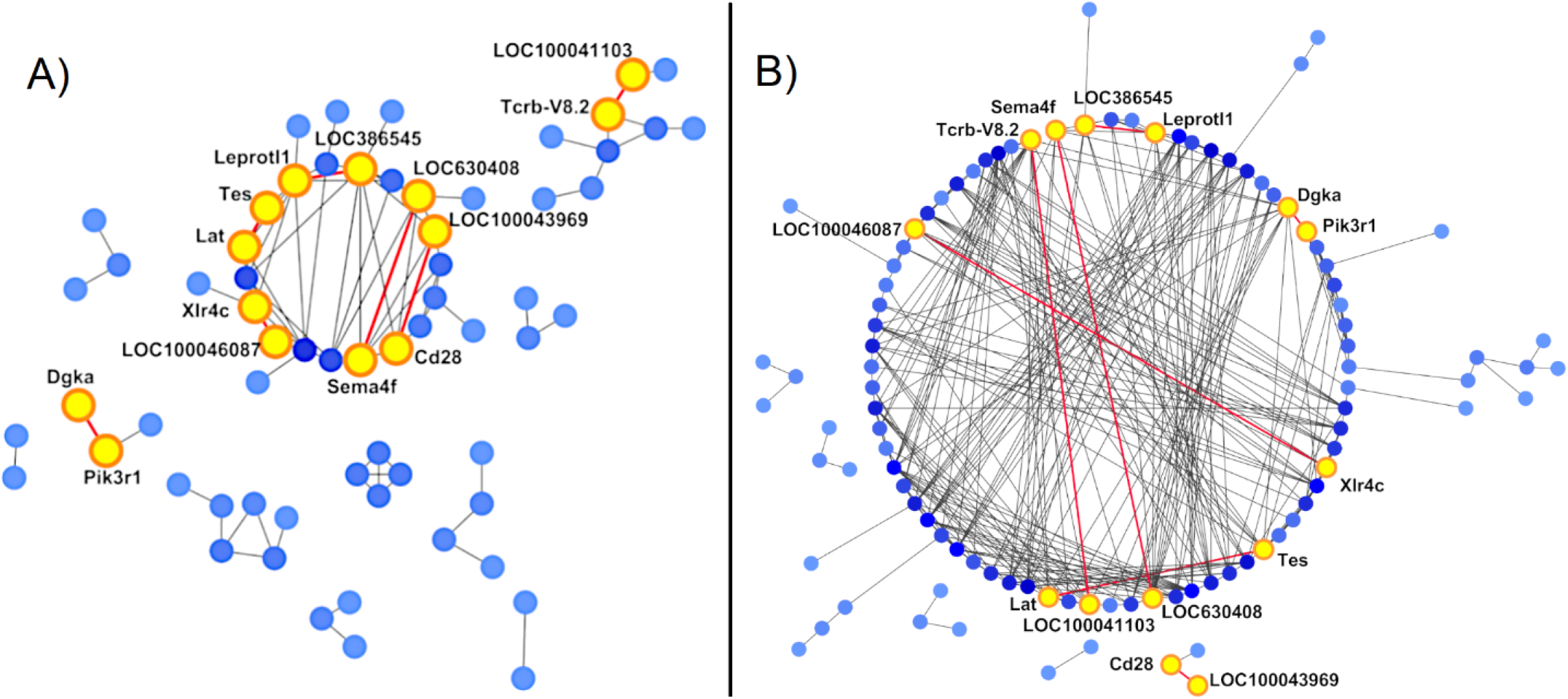
Network diagrams of gene co-expression with conserved edges between young and old splenic T-cells. The Spearman rank correlation reveals the gene co-expression network according to age. Darker colored nodes correspond with a higher number of edges per node (up to 10 edges in the younger mouse network and 15 edges in the older mouse network). A) The GCN constructed based upon gene co-expression in CD3+ splenocytes from 2-month old mice (64 nodes and 85 edges). B) The GCN constructed based on the old mouse data (102 nodes and 302 edges). The seven conserved edges between young (A) and old (B) mice are illustrated with the red lines. Node pairs at the ends of these edges are labeled and highlighted.

**Fig 4.**
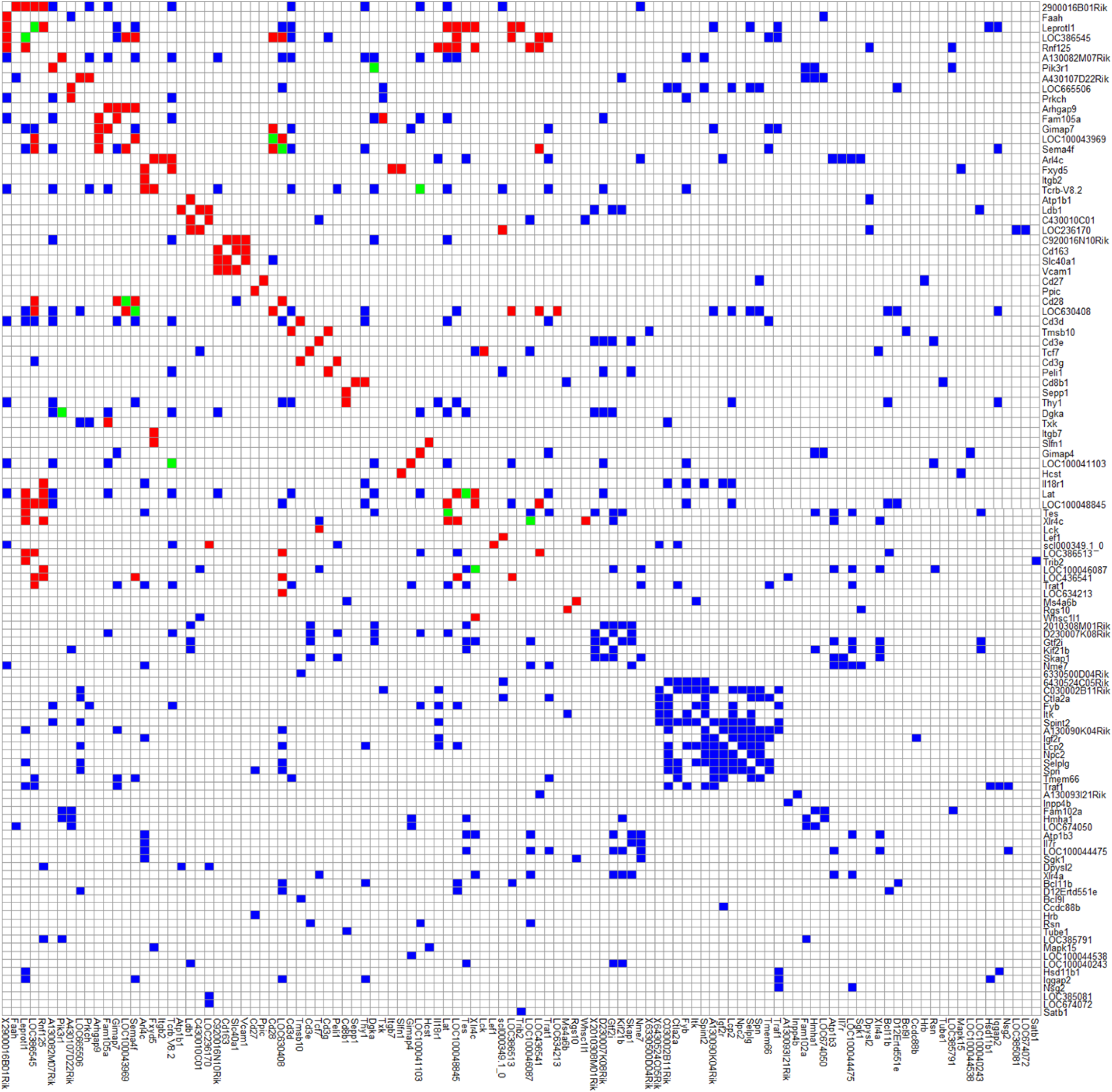
Adjacency matrices of gene co-expression for the two age related groups. The young adjacency matrix (red squares), the old adjacency matrix (blue squares) and elements that are conserved for both age-groups (green squares). Note that if you count the number of green squares above the diagonal, this is exactly seven which is how many conserved gene pairs are given in Table 3.

**Table 3:** List of the edges conserved between the young and old networks along with the genes (nodes) that the edges connect.

| Edge number | Edges Conserved Across Networks |
| --- | --- |
| 1 | Sema4f – LOC630408 |
| 2 | LOC386545 – Leprotl1 |
| 3 | Pik3r1 – Dgka |
| 4 | LOC100043969 – Cd28 |
| 5 | Tcrb-V8.2 – LOC100041103 |
| 6 | Tes – Lat |
| 7 | Xlr4c – LOC100046087 |

Consequentially, because only seven edges were conserved across the aging networks, most network edges were lost or gained with age. While the older mouse GCN saw an increase in total edges, 92% of the edges from the 2-month old GCN were not present in the 22/24-month old GCN, indicating 92% of the co-expression relationships established in young mice are absent in older mice. We identified 47 nodes present in the young network but whose co-expression relationships were lost with age in the senescent T-cells (Fig 5). Nodes in the older mouse GCN that gained new co-expression relationships not present in the younger mouse GCN can be viewed in Fig 6. Most of these genes with altered co-expression relationships encode for proteins involved in TCR/CD3 complex signaling and downstream signal transduction, such as the membrane receptor TCR alpha/beta chains, the CD3 complex (gamma, delta, epsilon CD3 chains), costimulation protein CD28, intracellular LCK, and Lat.

**Figure 5.**
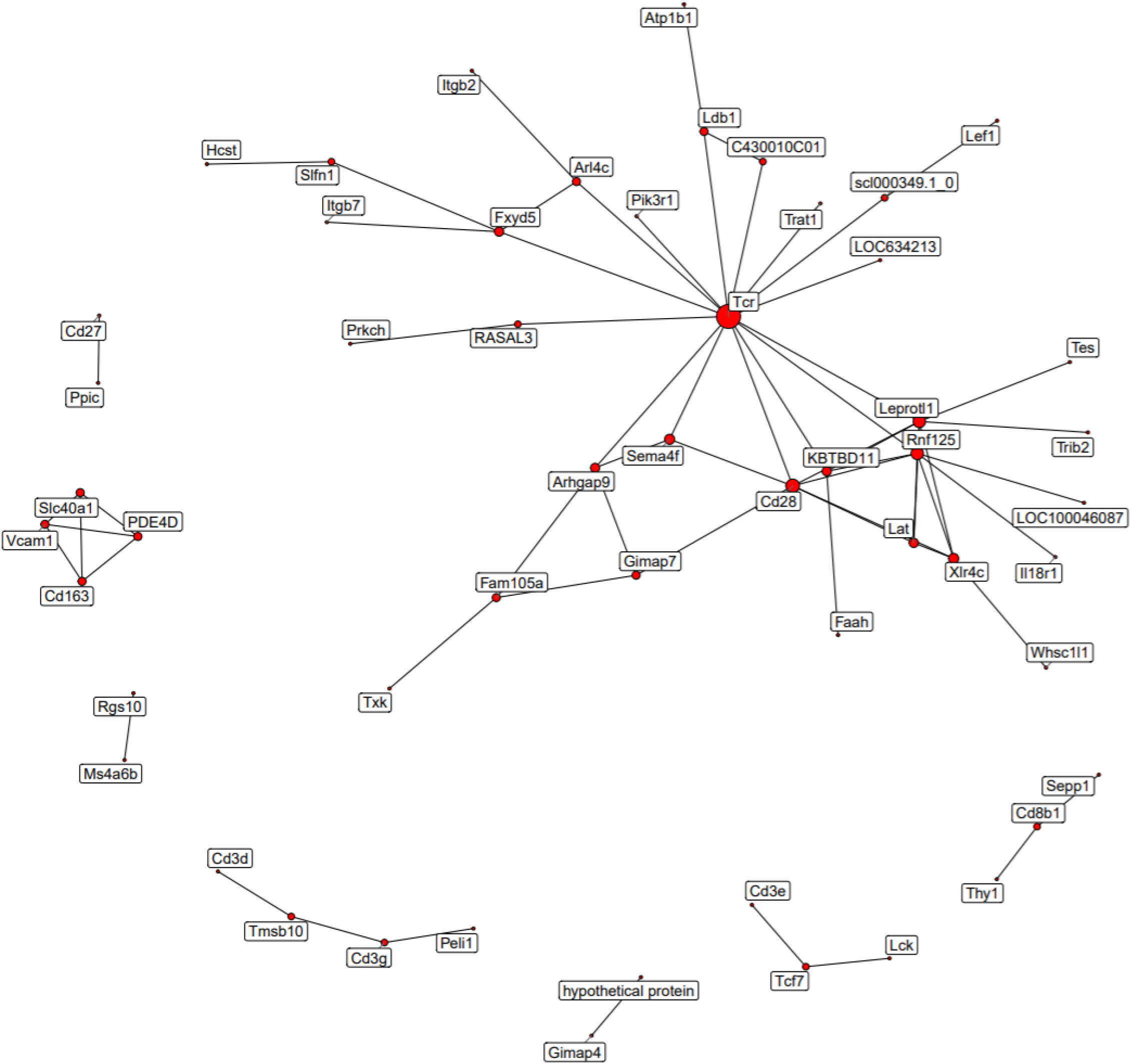
Genes with co-expression relationships lost with age. The young mouse GCN, containing only edges that are not in the 22/24-month old mouse network. TCR probes were merged into a single node to improve visibility.

**Figure 6.**
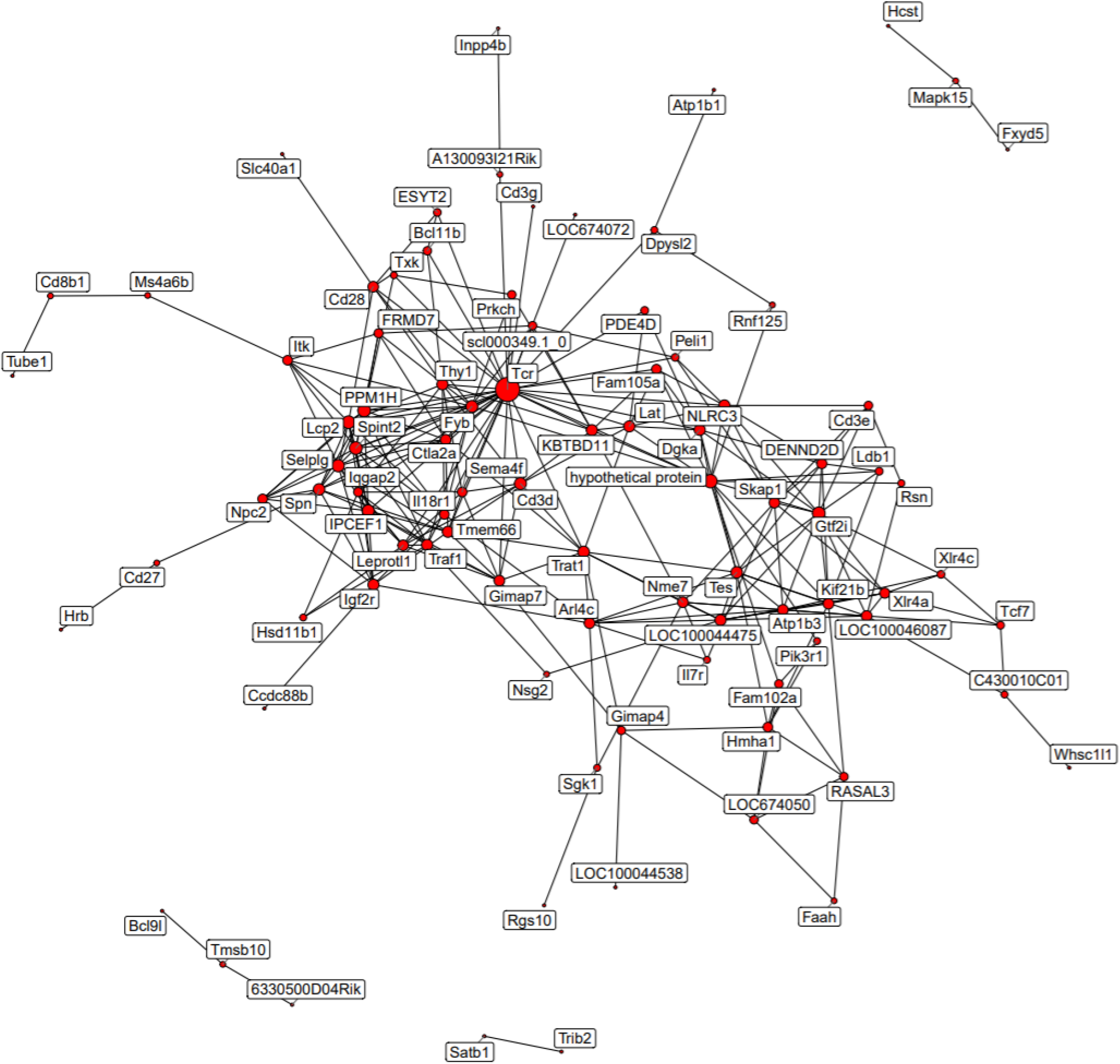
Genes with co-expression relationships gained with age. The older mouse GCN, containing only edges that are not also present in the 2-month old mouse network. TCR probes were merged into a single node to improve visibility.

### Network Centrality, Control and Node Power

In order to better quantify the evolution of topology of the network of T-cell gene expression during aging, we investigated the network centrality in the old and young mice network. The idea of network centrality has a long history [50]. In general, network centrality speaks to the impact of a given node within a given network. A number of network centrality variables have been subsequently formulated and discussed in the literature. More recently, these methods were applied to the study of longevity-gene networks [35–38] as well as in a variety of other areas such as neurological mapping and social network analysis [51–57]. We begin with the concept of “degree centrality”.

#### Degree centrality

*Degree centrality* counts how many neighbors a node has. A node has a neighbor if there is an edge directly connecting the node of interest to any other network nodes [35–38]. This definition argues that a network node is important if there are many nodes that are directly connected to it. From this definition, it follows that the given network node can affect the network to a greater extent, given that it has a high degree centrality. Degree centrality is the simplest of the centrality measures.

Table 4 illustrates the top ten gene-transcripts with the highest degree centrality in both networks. Of immediate note is the fact that the two young and old T-cell gene co-expression networks have totally different degree centrality nodes in the top 10 gene list, with the exception of TCR subunit gene LOC630408. Most genes with the highest centrality, including LOC630408, are identified as T-cell receptor components, which are highly diverse in the original spleen sample (S1 Table). In 2-month mice the degree centrality of TCR related genes is 6-10, while for the 22/24-month it doubles to 14 to 15, meaning that the TCR from senescent T-cells establish more links with other molecules. This suggests that the molecular relation is less specific (degeneracy), increasing the entropy of the system and decreasing the ordered organization established during development. Because degree centralities provide an overall view of the network connectivity, it is reasonable to hypothesize the fact that the 22/24-month old network is three times larger than the 2-month old network. We do note that network size differences may contribute to some of the overall differences in degree centrality differences.

**Table 4:**
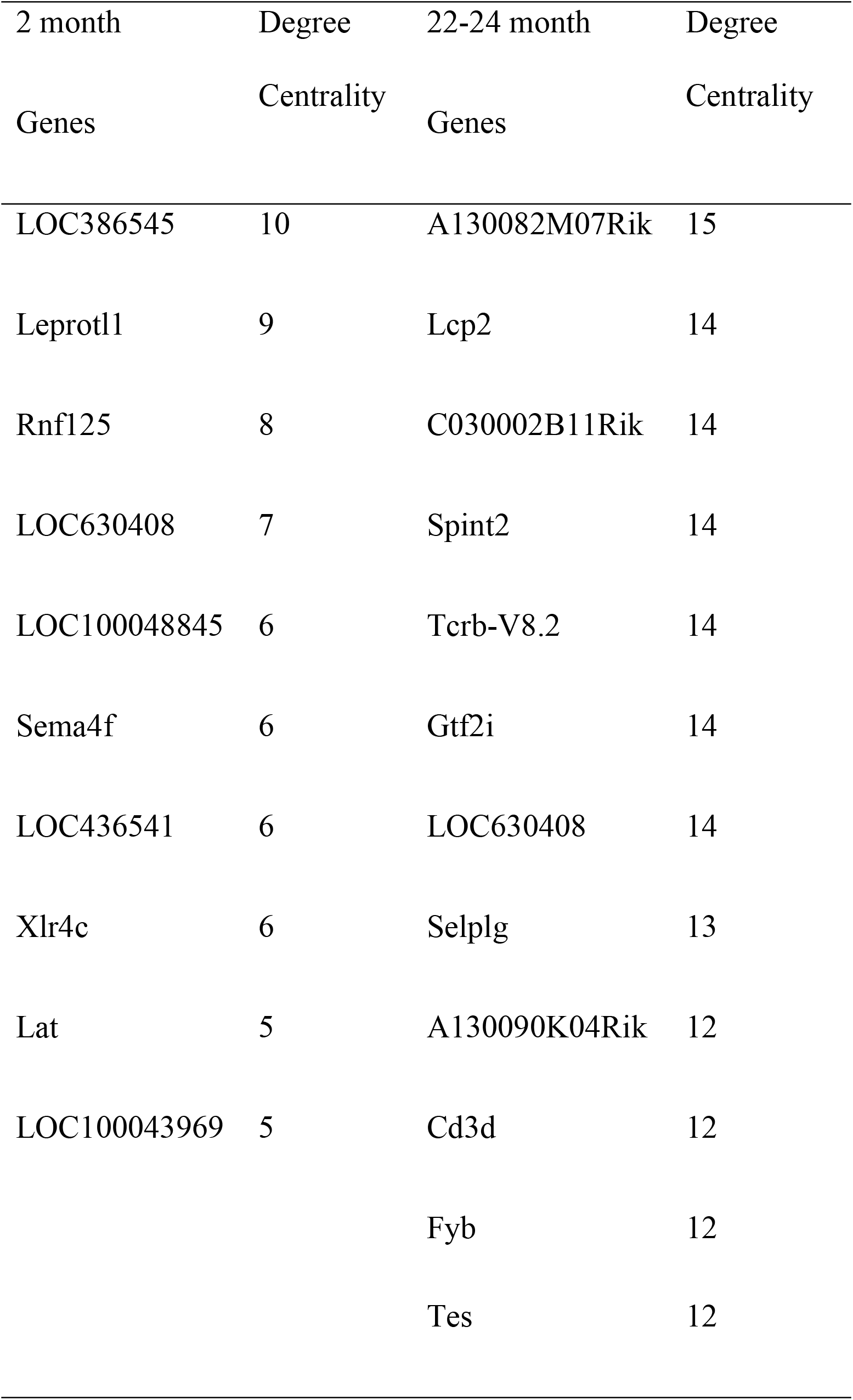
Top 10 highest degree centralities for the young and old mouse T-cell gene co- expression networks.

#### Eigenvector centrality

*Eigenvector centrality* is another measure of the influence of a given node in a network. A high eigenvector centrality means that the given node is connected to many other nodes who themselves have high eigenvector centrality scores. You can think of this as follows: if a node is a big shot, then its high eigenvalue centrality provides a measure of how many other big shots is it connected to.

Table 5 lists the top 10 network nodes with respect to their eigenvalue centrality value. These genes represent the top 10 genes that are thought to exert general control over the whole network. It is straightforward to observe that the two GCNs showed clear differences in hub control dynamics as defined by eigenvector centrality values. We note that the values of the 2-month eigenvalue centralities are all greater than those in the 22/24-month network, meaning a more connected and organized network. Additionally, we note that the genes that control the 2-month network according to eigenvector centrality are entirely different from the older mouse network, save for TCR gene LOC630408, which also has a high degree centrality in both networks as reported in Table 4. Most of the transcripts with high eigenvector values in young mice are involved in the TCR and the CD28 co-receptor. Meanwhile, the highest ranked gene in the old mouse network is Lcp2, which encodes the adaptor protein SLP-76 that is a central figure in the TCR signaling pathway. Information regarding the other genes with high eigenvector centralities can be found in S1 Table.

**Table 5.**
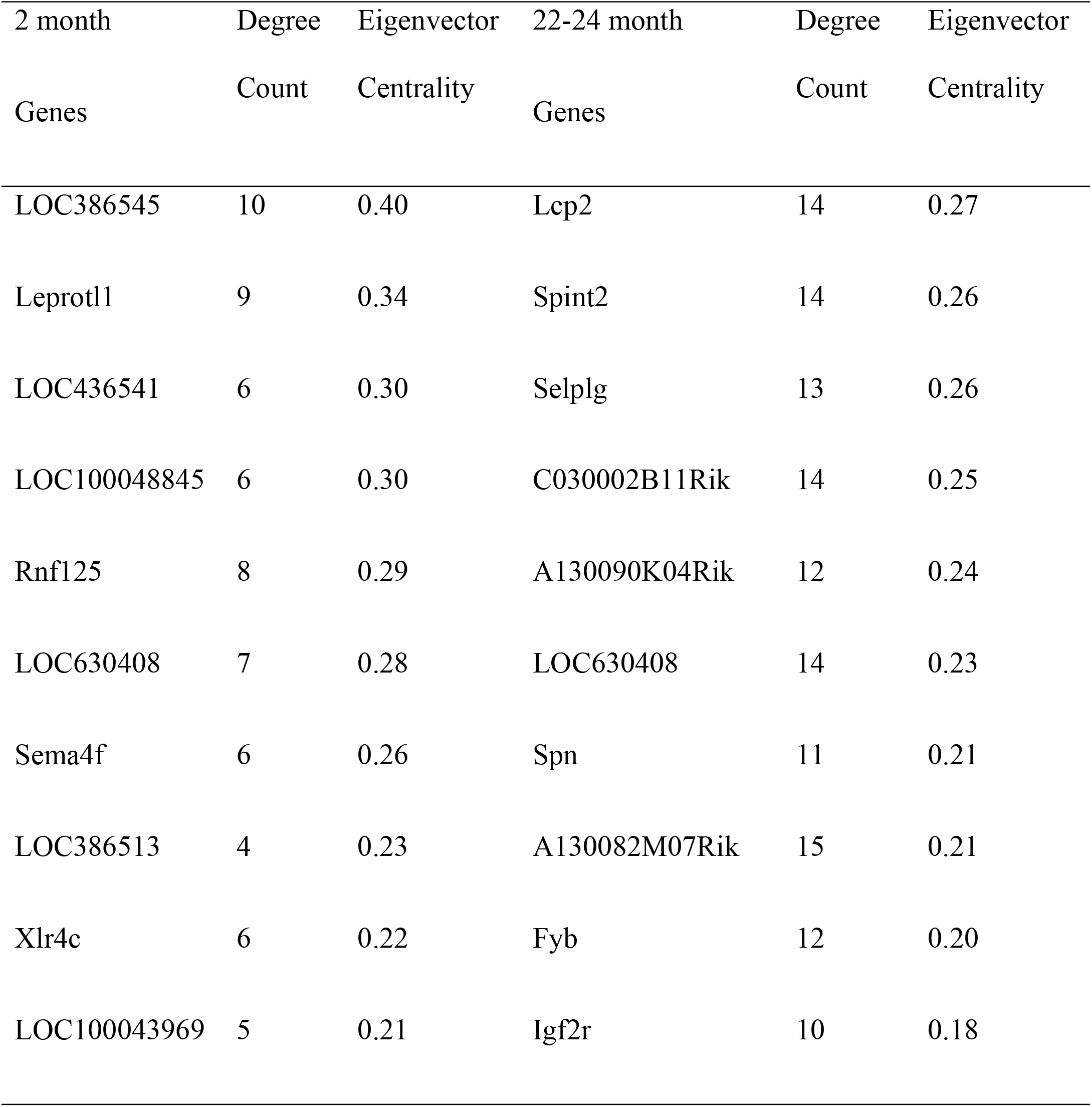
Top 10 highest eigenvector centralities for the young and old mouse networks.

#### Closeness centrality

*Closeness centrality* is a measure of the degree to which an individual node is near to all of the other nodes in a network. In Table 6 we provide the closeness centralities for the top 10 genes in both networks. First, we note that there are no common genes across the two networks. However, the functionalities of certain high ranking genes are conserved across the networks. For example, the first rank gene in the 2-month network is gene LOC386545 which is similar to T-cell receptor beta chain VNDNJC precursor (S1 Table).

**Table 6.** Top 10 highest closeness centralities for core the young and old mouse networks.

| 2 month Genes | Degree | Closeness | 22-24 month | Degree | Closeness |
| --- | --- | --- | --- | --- | --- |
|  | Count | Centrality | Genes | Count | Centrality |
| LOC386545 | 10 | 0.55 | A130082M07Rik | 15 | 0.38 |
| Leprotl1 | 9 | 0.51 | Trat1 | 11 | 0.38 |
| LOC436541 | 6 | 0.48 | Tcrb-V8.2 | 14 | 0.37 |
| LOC100048845 | 6 | 0.47 | Dgka | 11 | 0.37 |
| Sema4f | 6 | 0.46 | Lat | 11 | 0.36 |
| Rnf125 | 8 | 0.46 | Tes | 12 | 0.36 |
| LOC630408 | 7 | 0.45 | LOC100041103 | 10 | 0.35 |
| 2900016B01Rik | 4 | 0.45 | Cd3d | 12 | 0.35 |
| LOC386513 | 4 | 0.44 | Fyb | 12 | 0.35 |
| Cd28 | 5 | 0.43 | Arl4c | 9 | 0.35 |

Meanwhile first rank gene in the 22/24-month old network is A130082M07Rik, a gene that functions as the T-cell receptor alpha chain variable 9D-3 region (S1 Table). This change in the top-ranked closeness centrality hubs suggests differences in T-cell clonal expansions in the two age-related groups, as has been previously described [24]. The functions of other high-ranking genes can be found in S1 Table.

#### Betweenness centrality

*Betweenness centrality* represents the degree to which nodes stand in- between each other. One can say that betweenness centrality represents the control a specified node has over the network in that it finds nodes that act as bridges between nodes and – in a sense – these nodes may control the flow in the network. Such genes are often called bottleneck genes [58]. The higher the value of a node’s betweenness centrality, the greater the degree of control the specified node exerts over the network.

Table 7 lists the top ten gene betweenness centrality values for both networks. Betweenness centrality values for the 22/24-month network are significantly greater that those of the 2-month network. In fact, they are 5-10 times those of the 2-month network. The gene Rnf12 is common across both networks; although it does not have the same betweenness rank. Rnf125 is believed to function as a positive regulator in the TCR signaling pathway. In the 2-month network, the highest ranked gene is once more the TCR gene LOC386545, while diacylglycoprotein kinase α (Dgka) has the highest ranking in the older mouse network. Other high-ranking gene information can be located in S1 Table.

**Table 7.**
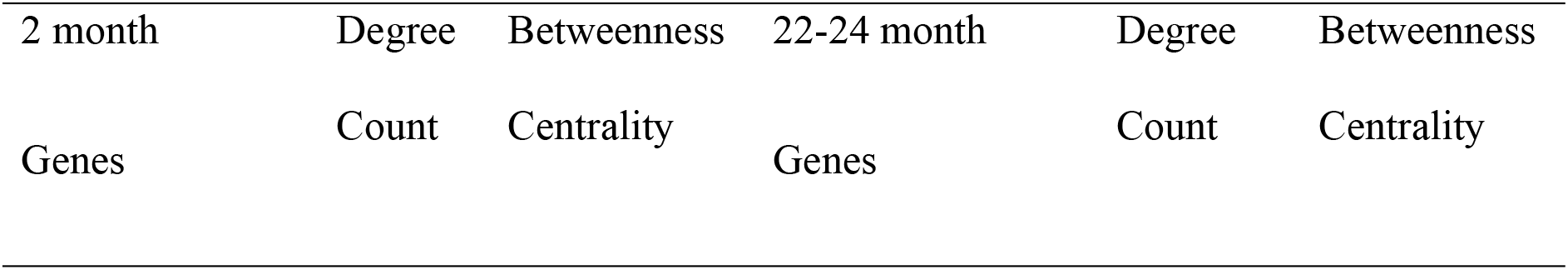
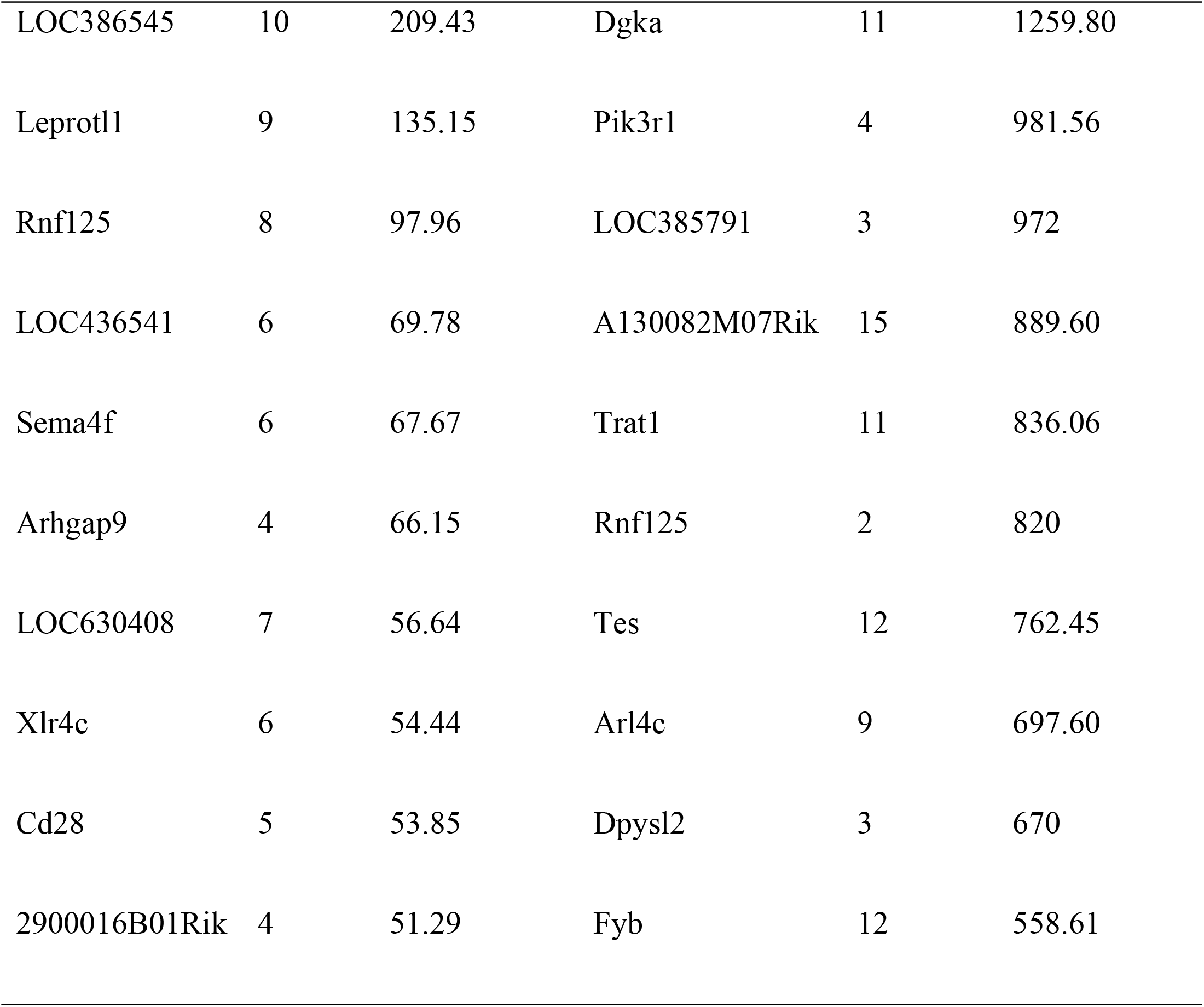
Top 10 highest betweenness centralities for the 2 and the 22-24 month old networks.

| 2 month | Degree | Betweenness | 22-24 month | Degree | Betweenness |
| --- | --- | --- | --- | --- | --- |
| Genes | Count | Centrality | Genes | Count | Centrality |
| LOC386545 | 10 | 209.43 | Dgka | 11 | 1259.80 |
| Leprotl1 | 9 | 135.15 | Pik3r1 | 4 | 981.56 |
| Rnf125 | 8 | 97.96 | LOC385791 | 3 | 972 |
| LOC436541 | 6 | 69.78 | A130082M07Rik | 15 | 889.60 |
| Sema4f | 6 | 67.67 | Trat1 | 11 | 836.06 |
| Arhgap9 | 4 | 66.15 | Rnf125 | 2 | 820 |
| LOC630408 | 7 | 56.64 | Tes | 12 | 762.45 |
| Xlr4c | 6 | 54.44 | Arl4c | 9 | 697.60 |
| Cd28 | 5 | 53.85 | Dpysl2 | 3 | 670 |
| 2900016B01Rik | 4 | 51.29 | Fyb | 12 | 558.61 |

#### Network Stress and Eccentricity

The maximum distance between a node and all other nodes is called the *eccentricity* of that node; biologically, this can be said to represent the easiness for all other genes in a network to reach another particular gene. Consequently, the higher the eccentricity value, the easier it is for a given gene to be influenced by the rest of the network – or, conversely, the easier it is for that node to influence the rest of the network [59]. The higher eccentricity in old mice may influence the time of processes and increase the delay in intracellular interaction pathways affecting efficiency of the cells. Table 8 lists the genes with the highest eccentricity values in both networks. We note that the eccentricity values in the 22/24- month network were higher when compared to the 2-month network. Whsc1l1 is the only gene with a highly ranked eccentricity value shared by both the young mouse and old mouse networks (Table 8). Whsc1l1 is defined as nuclear receptor binding SET domain protein 3 that methylates Histone 3 (S1 Table). The gene with the highest eccentricity in the 2-month network is Whsc1l1, and the highest ranked gene in the 22/24-month network is LOC674072 which is similar to Ig heavy chain V region 441 precursor.

**Table 8.**
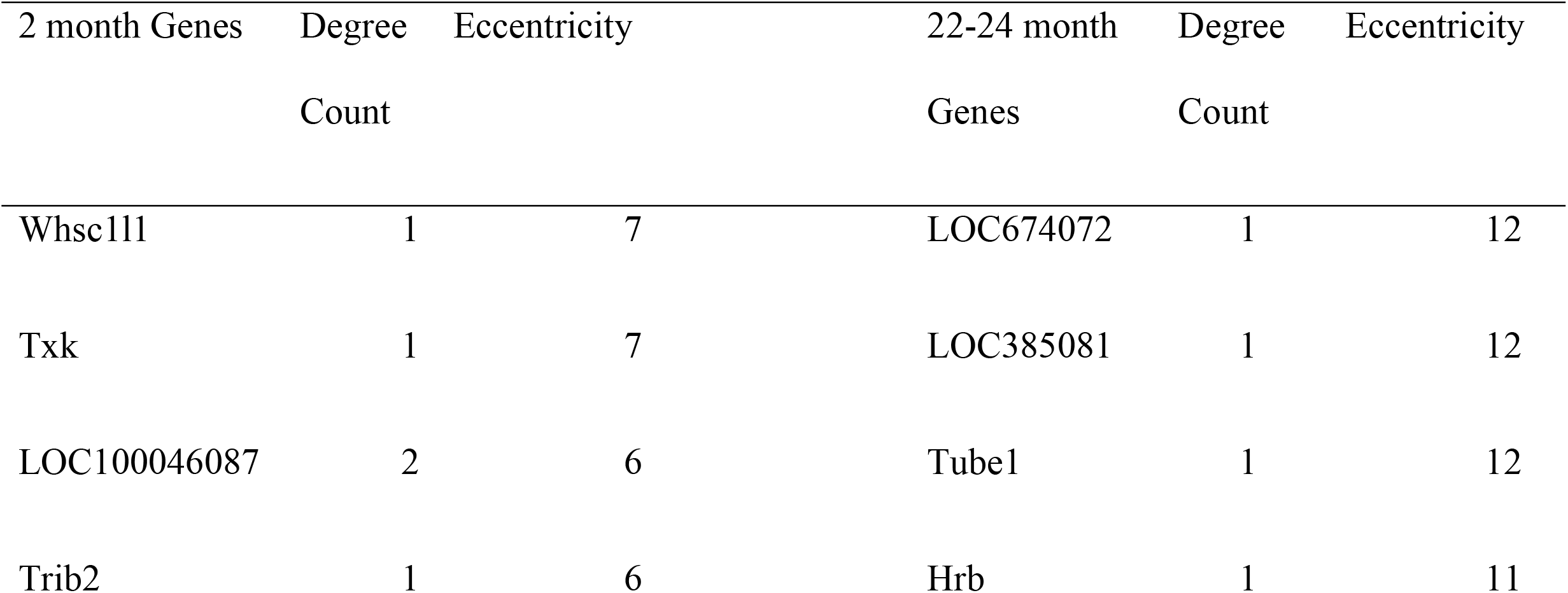
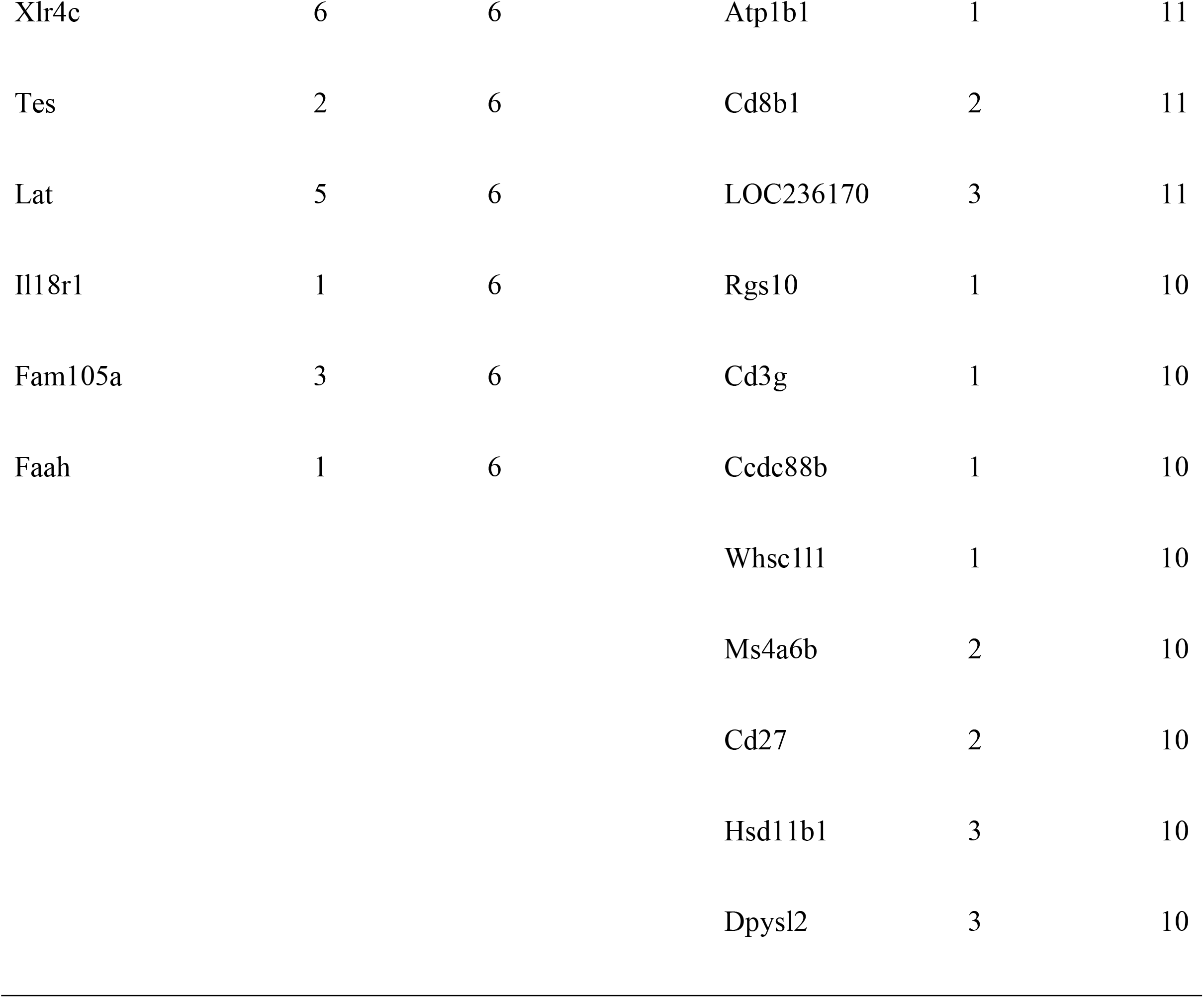
Top 10 highest eccentricity values for the young and old mouse networks.

*Stress* of a node in a biological network is defined as the number of shortest paths passing through a given node. It can be used to indicate the prominence of a gene in holding together connecting regulatory genes in a network pathway. The higher the value, the more importance the node has in holding together communicating nodes [59]. We find that stress in the top ten highest stress nodes in the 22/24-month old group appears much higher than the highest stress nodes in the 2-month old group (Table 9).

**Table 9.**
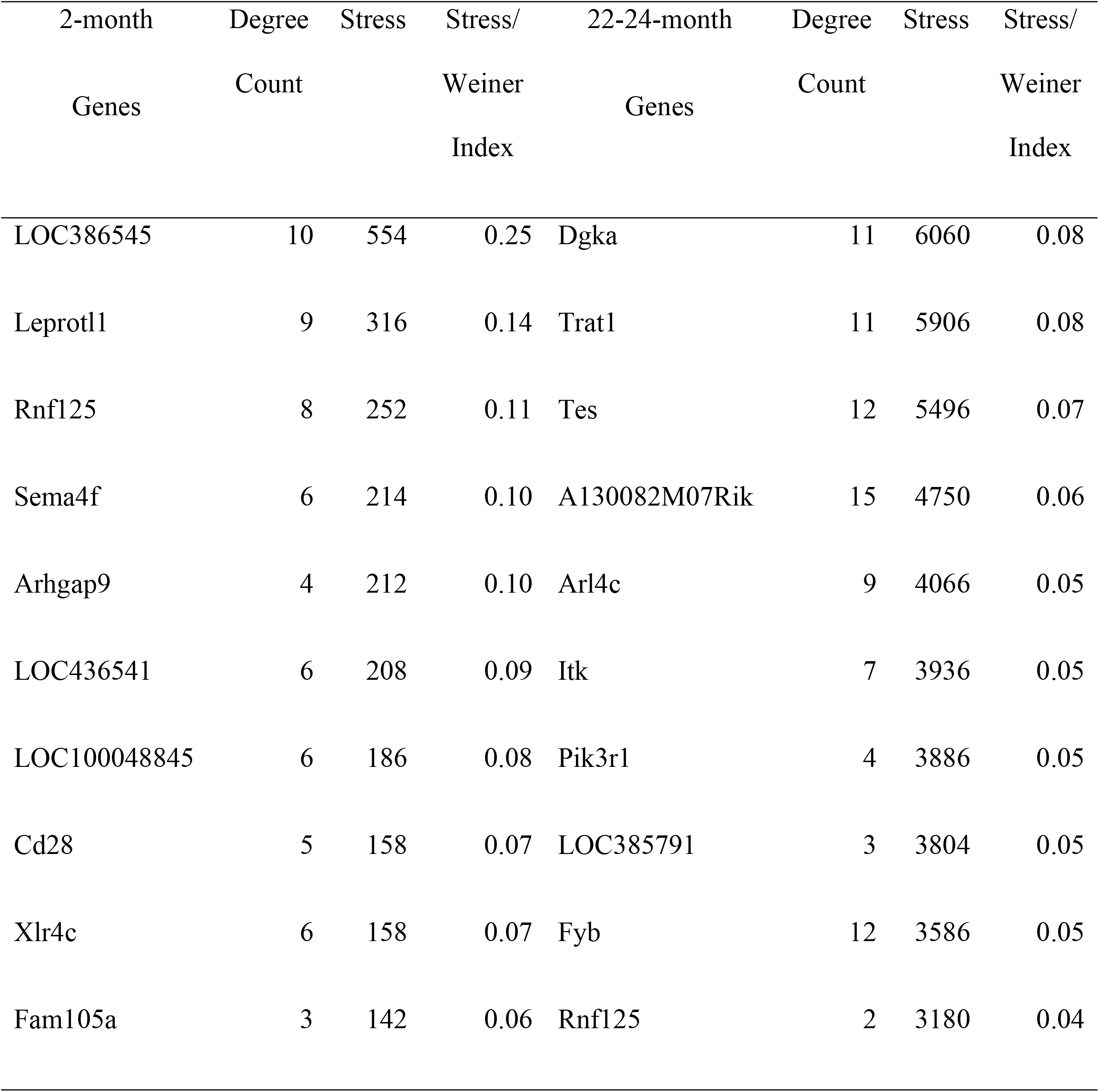
Top 10 highest stress values for the young and old mouse networks.

#### Weiner index

The *Weiner index* provides valuable context for these stress measures (Table 2). The Weiner index is the sum of the number of shortest paths in a network. We can use the Weiner index as a normalization factor. Comparing the ratio of the highest stress nodes to the Weiner index, we can see that the highest stress nodes in the older mouse group actually have a lower proportion of shortest paths in the total network as compared to the 2-month old group (Table 9). As a result, the genes with high stress values in the 2-month old group can be seen to have a greater impact on network connection than genes of high stress in the 22/24-month old group. Rnf125 is the only gene with high ranking in both networks and, as noted above, it is the only gene with high betweenness centralities shared by both networks as well. Rnf125 is similar to T-cell RNG activation protein 1. The function of genes with high-ranking stress values can be found in S1 Table.

#### Network Influence Measures

There are three *influence measures* of importance, where influence can be thought of as a measure of which nodes have power in a given network. These influence measures are given by: *(1) Local influence, (2) Indirect influence and (3) Total influence* [60]. The local influence of a node can be thought of as how much the given node affects its one-hop neighbors; the nodes directly connected to a given node by an edge. The indirect influence may be thought of as how much the given node affects its two-hop neighbors, nodes whose paths are connected by a single node mediating node. Lastly, the total measure of influence is defined as the weighted sum of the local and indirect influences. The ten highest local, indirect, and total influence values for the young and old mouse networks can be found in Tables 10-12. Again, a gene encoding TCR (LOC630408) is the only transcript in the top highest local, indirect, and total influence values across both networks. The identity of genes in the highest rank ordered influence values exclusive to each network can be found in S1 Table.

**Table 10.**
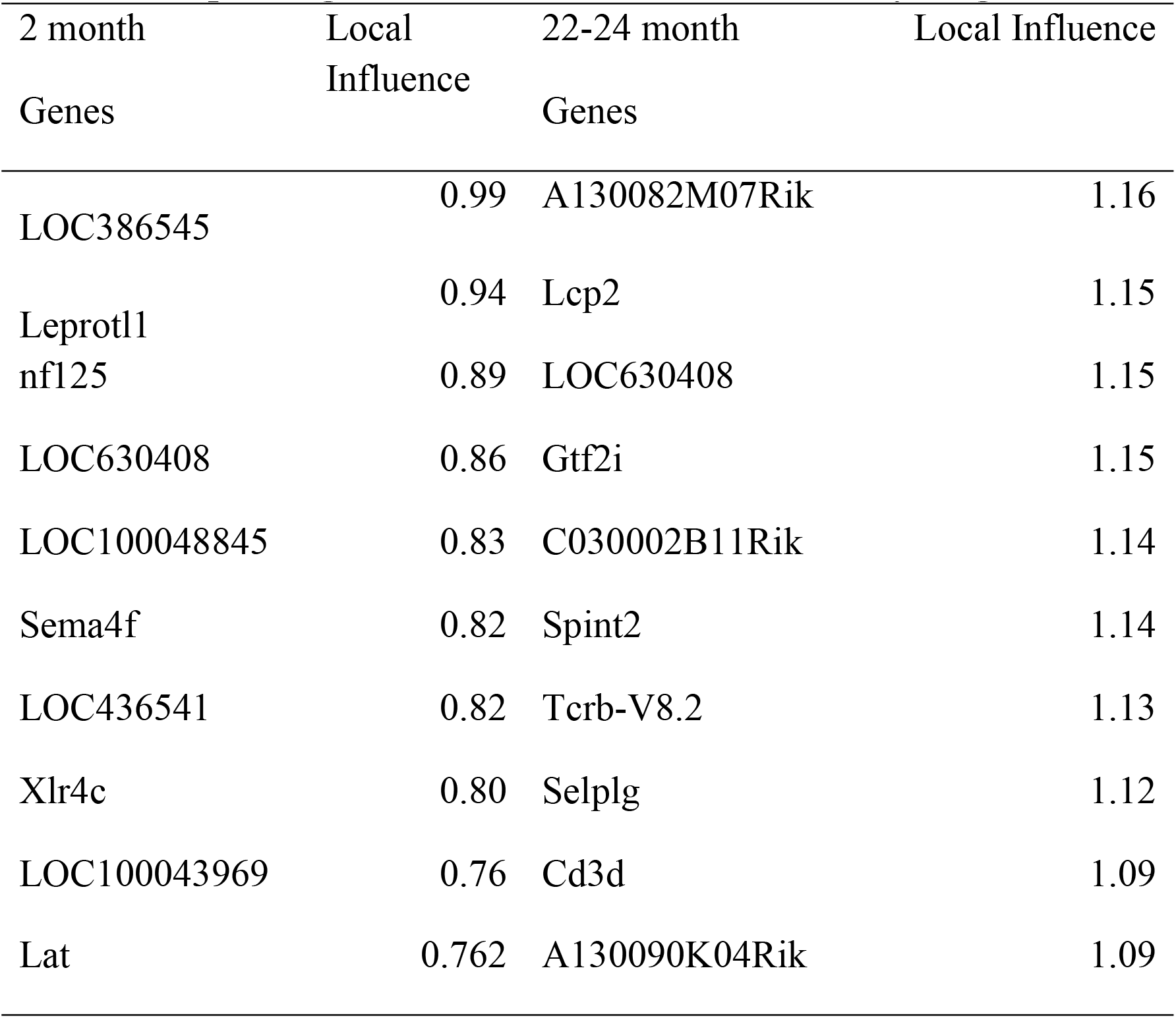
Top 10 highest local influence values for the young and old mouse networks.

**Table 11.**
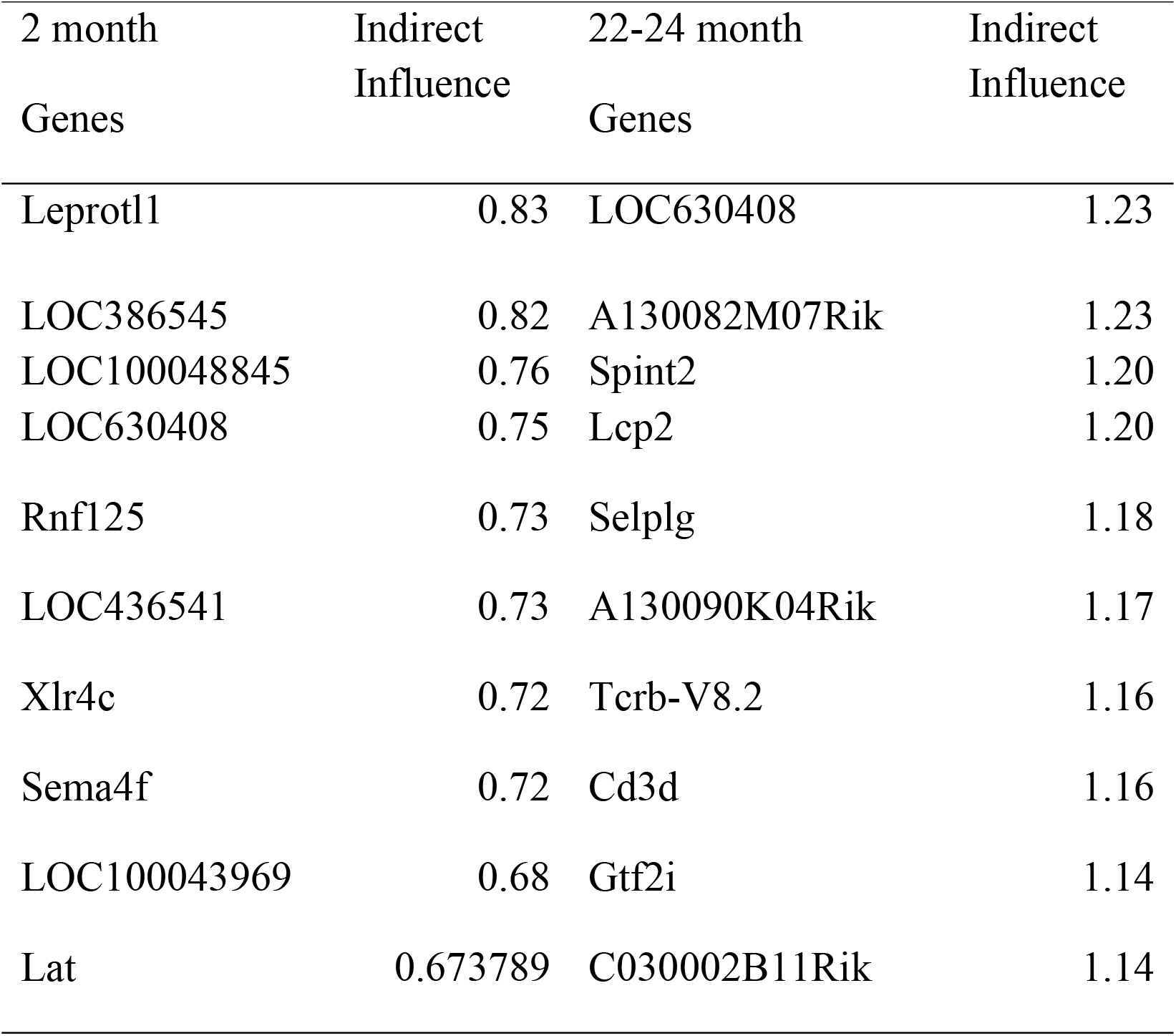
Top 10 highest indirect influence values for the young and old mouse networks.

**Table 12.**
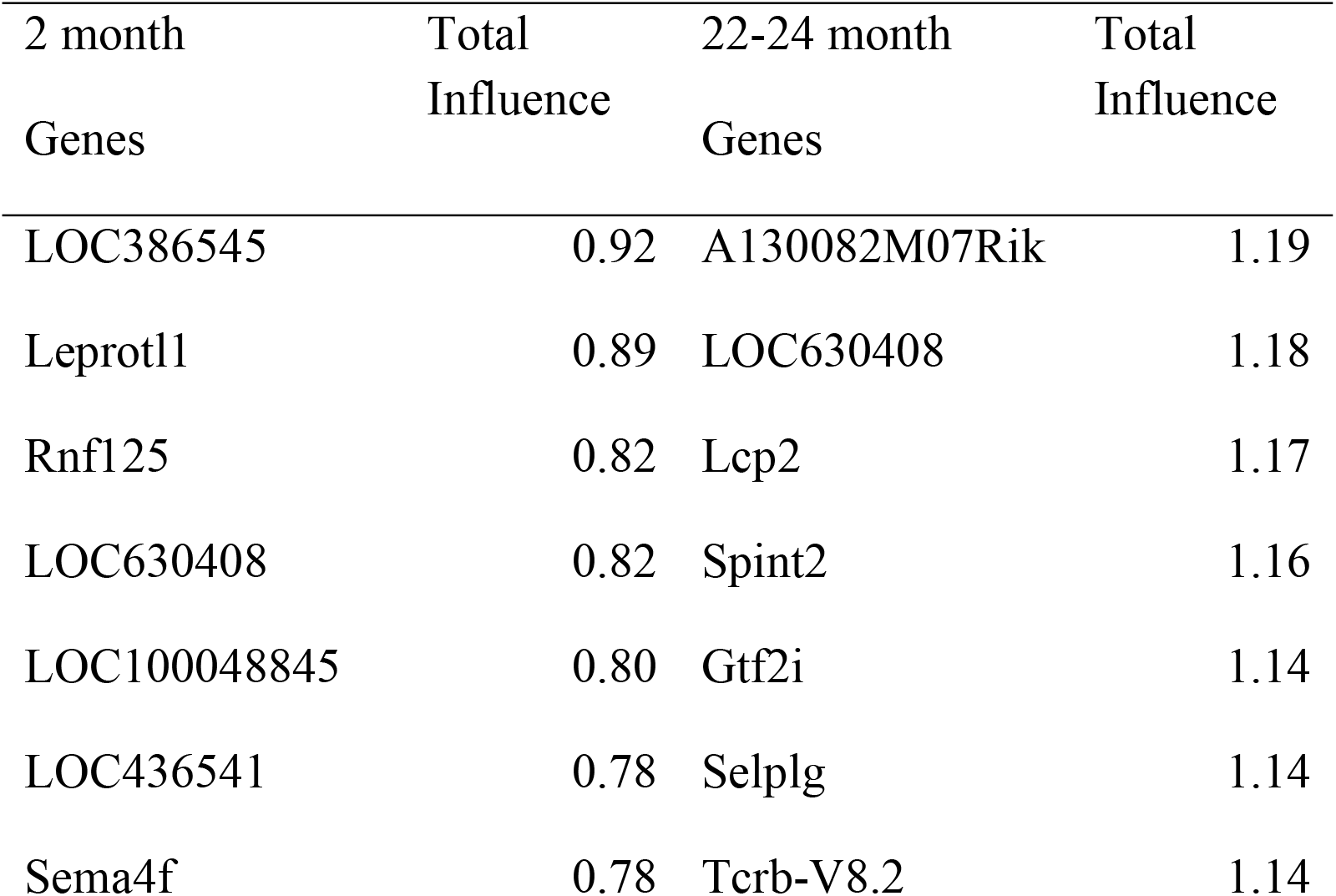
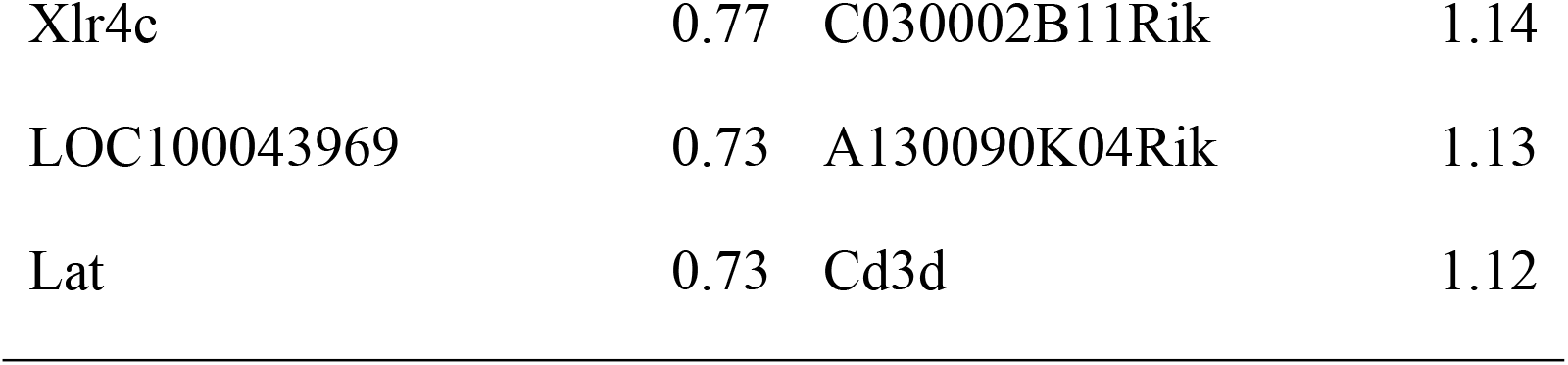
Top 10 highest total influence values for the young and old mouse networks.

#### Clustering Coefficient Analysis

The *clustering coefficient* of a node is a measure of the degree to which genes in a network tend to cluster together around the particular gene [28]. The mean clustering coefficient for each network, defined as the sum total of all clustering coefficients divided by the number of nodes, is found in Table 13. The mean clustering coefficient for the 2- month network is 0.251, while the mean for the 22/24-month is 0.335. The genes with the highest clustering coefficients in the 2-month and 24-month core networks can be found in Table 13. None of the genes with the highest clustering coefficients in the 2-month network were in the highest ranked genes for the 22/24-month network.

**Table 13.** Top 10 highest clustering coefficients for core the young and old mouse networks.

| 2 month | Degree | Clustering | 22-24 month | Degree | Clustering |
| --- | --- | --- | --- | --- | --- |
| Genes | Count | Coefficient | Genes | Count | Coefficient |
| Tes | 2 | 1.0 | C920016N10Rik | 4 | 1.0 |
| LOC100046087 | 2 | 1.0 | Hsd11b1 | 3 | 1.0 |
| LOC100043969 | 5 | 0.7 | Faah | 2 | 1.0 |
| Lat | 5 | 0.7 | Npc2 | 7 | 0.90 |
| LOC386513 | 4 | 0.67 | Fam105a | 7 | 0.86 |
| LOC100048845 | 6 | 0.6 | LOC100040243 | 4 | 0.83 |
| Sema4f | 6 | 0.6 | Ldb1 | 4 | 0.83 |
| Cd28 | 5 | 0.6 | Xlr4c | 5 | 0.8 |
| LOC630408 | 7 | 0.48 | Il7r | 3 | 0.67 |
| LOC436541 | 6 | 0.47 | D12Ertd551e | 4 | 0.67 |
|  |  |  | Tcf7 | 4 | 0.67 |

Fig 7 illustrates the spread of clustering coefficients *vs*. degree (k). Blue points indicate older mice, while red points indicate the younger mice. Of immediate note is that the 22/24-month network has a wider range of clustering coefficients compared to the 2-month network. Meanwhile, the 2-month network clustering coefficients are more compactly distributed. This apparent increase in highly interconnected genes in the old mouse GCN is consistent with the observed large core network that makes up the old mouse GCN (Fig 3).

**Fig 7.**
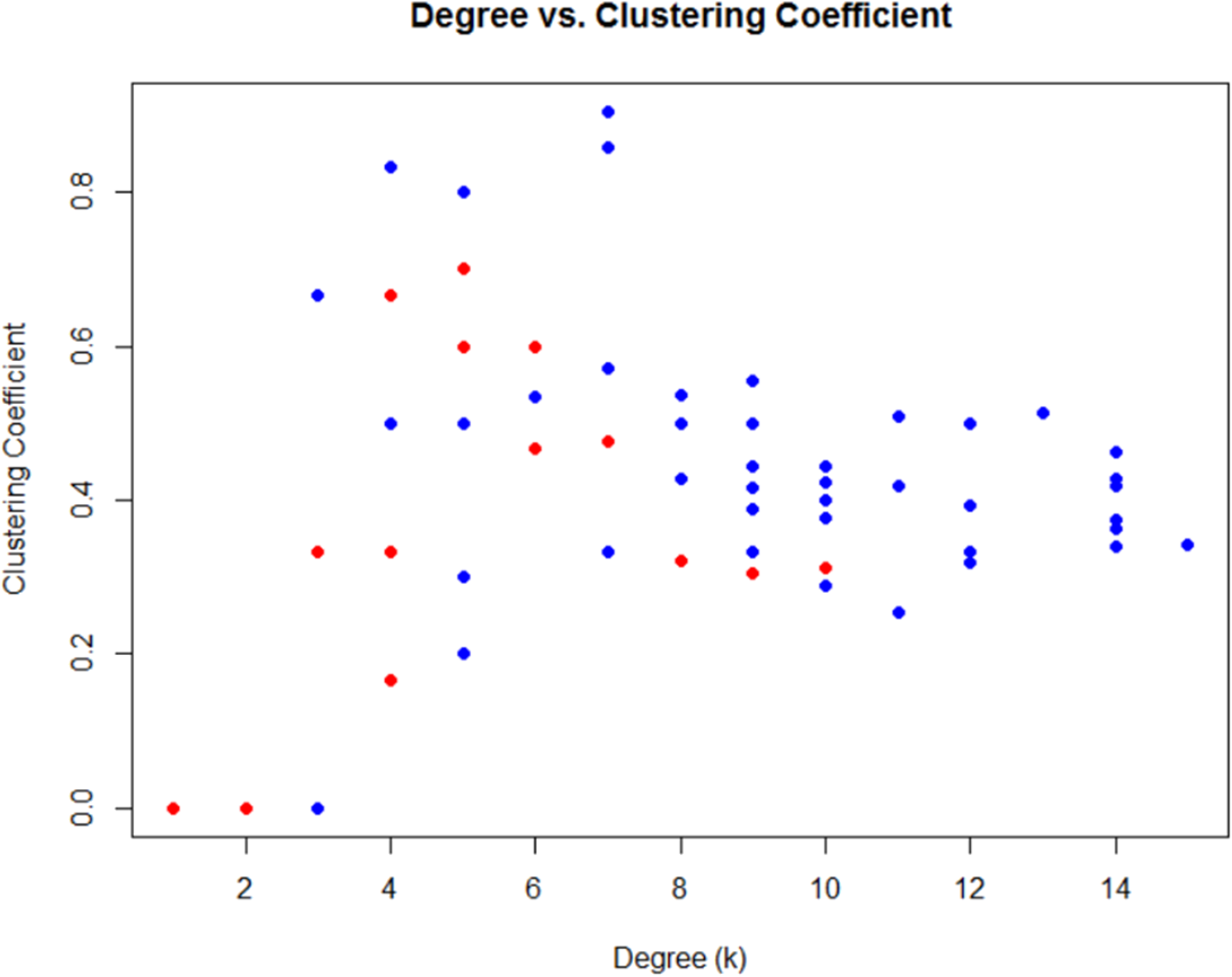
Plot of clustering coefficient versus network degree for the network gene co-expression in the two, mouse age-groups. Red points correspond with the 2-month age group, while blue corresponds with the 22/24-month age-group. It is straightforward to observe the large scattering of co-expression gene pairs in the 22/24-month age-group, while this is not the case for the 2- month old age group.

**Fig 8.**
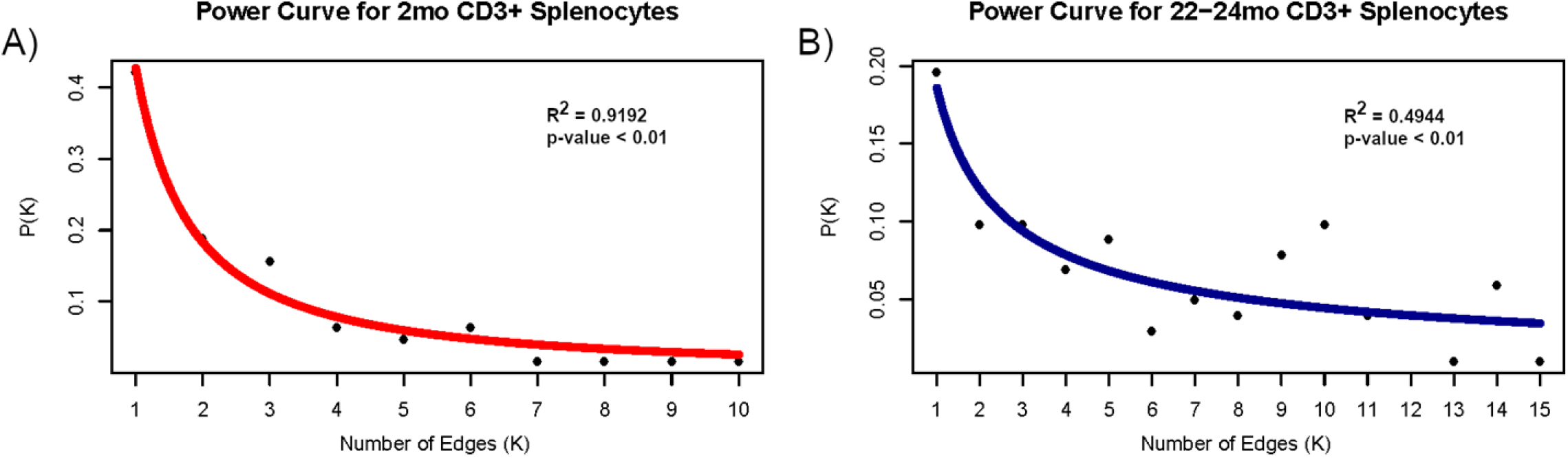
Power curve analysis of the two age-related networks based on degree distribution. Fitting the curve *P*(*k*) = *Ak*^―*γ*^: Probability of degree frequency “*k*” in (A) the two-month-old mice group can be moderately predicted by power law analysis (R^2^ =0.9192, p-value = 1.2x10^-5^, log P(k) = -1.58log(k) - 0.25). The predictive strength of this model was weakened in (B) the 22/24-month old group (R^2^= 0.4944, p-value=0.003452, log P(k) = -0.7461log(k) - 0.6883).

The program MCODE was used to find tightly connected subnetworks within a core network (http://apps.cytoscape.org/apps/mcode). MCODE is used to find clusters in network data based upon core clustering coefficients. It can determine pathways of biological significance in protein and gene networks. MCODE found four tightly linked subnetwork clusters in the 2-month old network, and six tightly linked clusters in the 22/24-month old network (S3 Fig). The identity of genes involved in these subnetworks can be found in S1 Table.

#### Centrality Analysis of Conserved Gene Pairs

Finally, we summarize the rankings of the conserved co-expressed gene pairs (see Table 3) in Table 14. We note that Leprotl1 and the TCR gene LOC386545 have most of the highest rankings among the genes that are conserved between the young and the old networks.

**Table 14.**
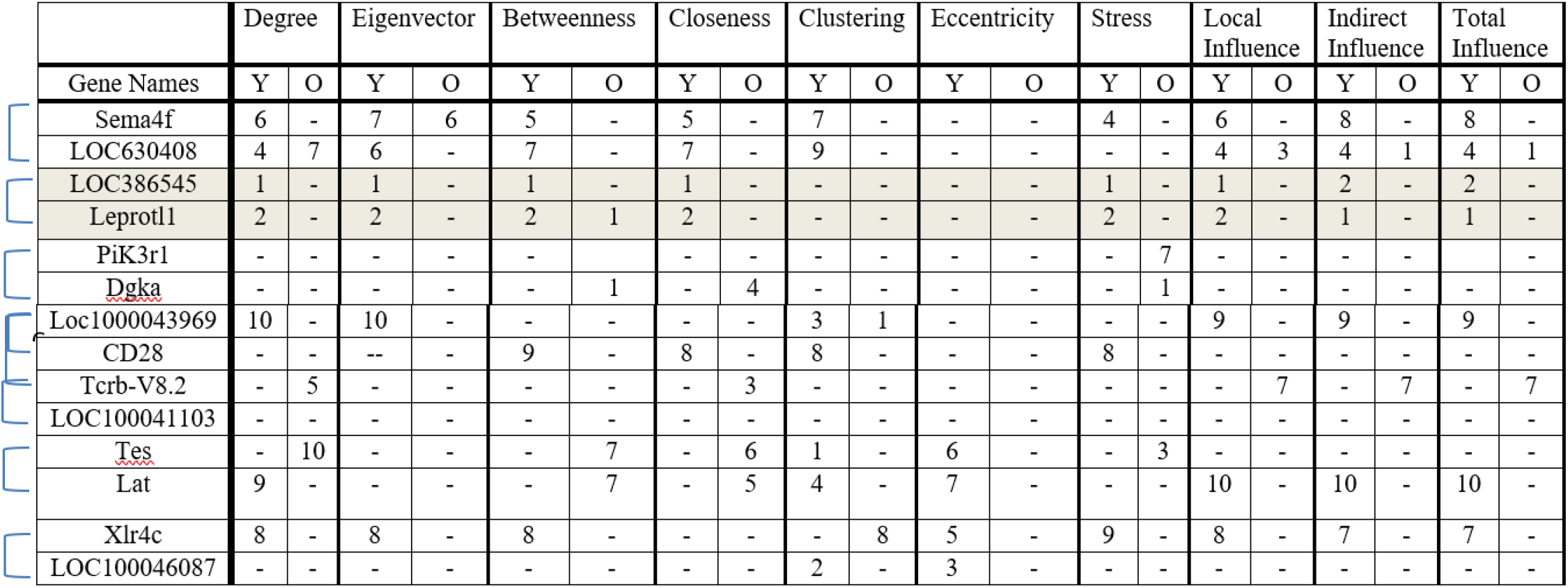
Centrality rankings of genes with conserved co-expression in both networks.

Column headers indicate the age of the mice from which the samples were taken (Y: 2-month, O: 22/24 month). A dash indicates that the given gene was not in the top 10 genes for that centrality variable. The braces correspond to the nodes (genes) at the end of the edges in Table 3. We note that the edge with the gene pair LOC386545 (similar to T-cell receptor beta chain VNDNJC precursor) – Leprotl1 (Leptin receptor overlapping transcript like 1) appears across all centrality measures, except for clustering and eccentricity, in that it always ranks 1-2 for the young mouse cells but not for the old mouse cells. This makes this edge and nodes pair a very powerful component in the young network.

### Scale Free Analysis

Scale-free analysis has significance in biological networks [61–63]. Scale-free networks are networks whose degree distribution follows a power law with a power parameter that typically falls in the interval 2-3, however, it may fall outside this interval [64]. Analysis of our two network architectures showed that the two-month old GCN is of scale free type, while the scale-free network structure was weaker for the older mouse GCN. Power curve analysis, fitting the function *P*(*k*) = *Ak*^*γ*^ to the P(k) for each network, showed that the younger mouse GCN followed a strong power law behavior (R^2^ = 0.9192, p-value < 0.001, Fig XA), while the older mouse group demonstrated a weaker power law behavior (R^2^ = 0.4944, p-value <0.01, Fig XB).

## Discussion

The gene co-expression network analysis of a 130 gene signature from young and old mice yielded a variety of interesting insights and conclusions about aging. First, we observe an alteration of the TCR signaling pathway in old mice, where the TCR complex and CD28 co-stimulatory molecules upstream the pathways are particularly affected. The RNA encoding the proteins present on the membrane of T-cells required for full T cell activation were co-expressed in the young mouse GCN, while this was not true of the old mouse network. The small core network of TCR-related genes in the young mice reveals that gene co-expression for proteins involved in TCR signaling are stable across the three strains of mice, suggesting the core network is robust across the genetic variation in mice strains.

Further, while the network of young mice is well organized and reflects controlled development across each genetic background, those of old mice appears more stochastic. We found a 255% increase in the number of edges in the 22/24-month old network when compared to the 2- month old network. However, we only found seven edges in the 22/24-month network that were also found in the 2-month network. This suggests that many of the edges in the 22/24-month old GCN were the result of novel co-expression relationships likely gained as a result of age.

Power law analysis suggests a deterioration in network organization in the old mouse network compared to the young mice. Analysis by power law demonstrated the 2-month old network was a scale-free network type, while the 22/24-month old network was not. Scale-free networks are frequently more robust to perturbations than are other network architectures, indicative of both complexity and organization. The scale-free network structure of the 2-month GCN is typical of most biological networks [62]. Scale-free network types may provide an advantage to biological systems that are subject to damage by mutation and/or viral infection. The scale-free network structure of the young CD3+ T-cells indicates that transcription of genes is tightly regulated in the splenocytes of 2-month old mice that are fully immunocompetent. Decrease in scale-free structure in the co-expression patterns of the 22/24-month old mouse splenocytes suggests that this control is decreased with age.

This change in network architecture has interesting implications for the regulatory behavior of genes in older T-cells. Because the major signal transduction from the TCR complex and CD28 is disturbed, the expression of the internal cell molecules is consequently perturbed in senescent T-cells. That the 22/24-month old mouse splenocytes see an increase in correlation at the expense of more organized network structures suggests T-cell regulation and immune behavior is diminished. Indeed, in the same series of FVB and B6 mice, we previously quantified by flow cytometry and mathematical models that the T-cell cycle transition rates and proliferation/death rates were altered in 18 month-old mice as compared to 2 month-old mice [24]. And while we have no direct evidence for the implication of these results with respect to human immune T-cell dynamics, we do know that human T-cell populations are also altered with age, and that there are a variety of other changes in T-cell behaviors in humans: in particular, changes in signal transduction in T-cells, and the promotion of long lived T-cells with less efficient function and inflamm-aging [65], even while the total lymphocyte maintenance is preserved [66].

Changes in network structure with age also reveal differential co-expression relationships in several key gene pathways. For example, expression of Transcription Factor 7 (TCF7) is important for T-lymphocyte differentiation. This transcription factor is essential for the proliferation of early DN thymocytes, the survival of DP cells, positive/negative selection, as well as for the creation of the CD3/TCR complex and the cell cycle control, and, in peripheral tissues, for the orientation to the TH2 pro-inflammatory pathways and activation of memory CD8 T-cells [67]. We previously quantified increased time spent in gap phases of the cell cycle leading to decreased division rates in immature DN and mature CD3hi DP thymocytes in FVB mice: the time spent in gap phases in G0/G1 and G2/M2 from DN to CD3hi DP increases from 10.2 days in young B6 mice to 28 days in young FVB mice, and even up to 58 days in 18 months old FVB mice. These proliferation rates were decreased in thymus and the spleen in FVB mice, as compared to B6 mice, contributing to acceleration of the thymic involution [24]. We also found increased occurrence of oligoclonal expansions of CD8 T-cells in FVB old mice [25].

Additionally, we previously observed an increase in differentiation of Tregs in old FVB mice but defective proliferation in the spleen. Interestingly, such a defect could be related to a deficiency of TCF7 in old FVB mice. Such a defect would lower the threshold to differentiate Tregs.

In the 2-month old network, TCF7 falls in a three-node island with genes Lck and CD28e. Transcription of TCF7 is previously reported to correlate with the expression of both Lck and CD28e [68–69], and the 2-month old network indicates these biological relationships. This co-expression linkage is lost in the 22/24-month old GCN, where the TCF7 node loses connections to Lck and CD28e and gains co-expression relationships with genes Xlr4a, Xlr4c, C430010C01 and TCR probe LOC10004608. This suggests massive alteration in this gene regulatory pathway with age. Because TCF7 increases Th2 differentiation and inflammation, these changes in co-expression relationships might contribute to inflammation [67], in addition to changes in T-cell populations observed with age [24] [25].

However, it is not fully known if the increase in co-expression relationships across the old mouse GCN serves any biological function, or whether these changes have a detrimental or compensatory impact on gene expression behavior in old T-cells. Some results in the original signature, however, suggested biological relevance with what is currently known about T-cells during immunosenescence. The old mice had lower expression of these genes related to TCR activation, such as CD3, VB8.2, co-stimulation molecule CD28, early activation molecule CD69, adhesion molecule VCAM1, Thy1, receptor to lymphokines IL-7R, and CD27. The TCR/CD3 complex receptor-to-antigen interaction, and the downstream signal transduction involving LCP2/SLP-76 recruitment, binding GADs and lat, the co-stimulatory signal via CD28 and its cytoplasmic tail binding, and CTLA-4 are all affected by age [49–50]. We observed that the aging signature affects several functions related to hematopoietical system as seen in IPA (S2 Fig), related not only to activation (TCR/CD3/CD69), but also binding and adhesion/interaction (VCAM1). Pathway enrichments for the genes appearing in both networks also suggest these genes play a role in changes in the immune system with age (Fig 2).

Differences in network hubs within the two age-related networks can be used to infer more about changes in key gene pathways that may be attributable to age. Hubs are important determinants of system functions in a biological network; in co-expression networks, deletion of a hub gene is likely to result in cell death [70]. Across the two age-related GCNs, very few of the hub genes as defined by degree, eigenvector, betweenness, and closeness centralities, as well as the influence measures, clustering coefficient, eccentricity, and stress values, were the same (Tables 4-13). Further, very few hub genes in the older mouse network were highly rated across the different node centrality measures, while there was more consistency and less variety of highly-rated genes in the young mouse network. This suggests the hubs in the young mouse network have overall more control, importance, and power over the network structure compared to the old moue network hubs.

Two genes with a co-expression relationship conserved across both networks, Sema4f (sema domain, immunoglobulin domain (Ig), TM domain, and short cytoplasmic domain) and TCR (gene LOC630408, T-cell receptor alpha chain V region CTL-L17 precursor), appeared in a high rank across multiple hub measures for both age-related networks, as summarized in Table 14. Additionally, both Sema4f and TCR/LOC630408 were incorporated into a tightly clustered MCODE subnetwork for the young mouse group (S3A Fig), but not the old mouse network (S3B Fig). That the co-expression relationship between these genes is conserved in both networks implies this expression relationship has important biological function, or a precursor to these genes serves as an important biological mediator. Further, that these genes serve as important hubs in both networks suggests a biological importance of both genes in both young and old splenocytes, although the lower centrality rankings and lack of MCODE clustering for the older mice suggests these genes have a diminishing role with age.

We note that the edge of co-expression with the gene pair TCR (LOC386545 similar to T-cell receptor beta chain VNDNJC precursor) – Leprotl1 (Leptin receptor overlapping transcript like 1) appears across all centrality measures, except for clustering and eccentricity, in that it always ranks 1-2 for the young mouse cells but not for the old mouse cells (Table 14). This makes this edge and nodes pair a very powerful component in the young network. The Leprotl1 gene encodes a homologue of the leptin membrane receptor. Leptin is an adipokine than regulates appetite and food intake but also is involved in T-cell differentiation and regulation of adiposis and the immune system. Notably, the exponential rise in metabolic and autoimmune diseases with age might be related to metabolism and leptin. Leptin is secreted by both adipocytes and T-cells, in particular this autocrine production by Treg cells can down-regulate T-cell proliferation during immune response and prevent inflammation [71]. Interestingly, leptin inhibits the proliferation of Treg cells [72] and their control of inflamm-aging. The leptin receptor is further involved in JACK2 STAT-3 orientation and Thelper cell activation, promoting Th17 differentiation but inhibiting Treg differentiation. Moreover, leptin promotes the proliferation of naïve T-cells but inhibits the proliferation of memory T-cells. The Leprotl1 gene is highly expressed in FVB mice, as is CD69, an early activation marker of T-cells. Notably, by contrast, these proteins are lesser expressed in B6 and BALB/c mice. The conservation of co-expression of Leprotl1 and the TCR gene across the age-related networks stresses the biological importance of this co-expression relationship, but changes in network hub rankings suggests the downstream pathways of the leptin receptor are altered with age.

In conclusion, we found significant deterioration of network organization when comparing the 2-month network to the 22/24-month network and observed changes in hub genes suggesting changes in TCR signaling and co-stimulation through CD28 in the two age-related groups. This analysis provides evidence of a defective organization of transcription in older peripheral T-cells. This dis-organization is suspected to increase delays in T-cell regulation pathways and inhibit the biological activity of CD3+ lymphocytes, while other cells die and cannot be quantified here. In addition, the identification of hub genes for network expression in the young and old mouse groups help identify genes important to healthy cell function and deteriorating function with age, providing opportunities to approach these genes as potential therapeutic targets to help aging patients. Human orthologs of the genes used in this analysis (S1 Table) could help to understand human aging in lymphocytes.

### Limitations

Previous GCN research using AGEMAP data found a decrease in gene co-expression with age in mice across multiple tissue types [38]. This contrasts with the results of this study, which found a nearly 255% increase in the number of edges in the old mouse group GCN as compared to the younger mouse group. There are several factors that may contribute to this result. One component may be experiment design: the AGEMAP study utilized tissue from only C57BL/6 mice, both male and female, instead of using exclusively female mice from three separate mouse lines. Additionally, samples in the AGEMAP study were extracted from mice at ages 16 and 24 months, as opposed to 2 and 22-24 months. The size of both datasets may also contribute in different, limited results (10 mice per age/sex, as opposed to 11-12 mice per age group). Finally, an important factor is that the AGEMAP study did not look at co-expression in peripheral T-cells, which may behave differently from other cell types across the aging process. Indeed, T-cells are permanently selected to survive and to divide through the TCR complex and co-stimulation pathways, or they undergo apoptosis if too many default signals occur (as during the cell cycle). Results by Vibert & Thomas-Vaslin [24] show that across ages and genetic backgrounds, immature T-cells in the thymus and mature T-cells in spleen show a decrease in T-cell proliferation with early T-cell differentiation, compensated by an increase of proliferation of effector/memory T-cells in spleen, that often correlates with oligoclonal T-cell expansions [25]. Because 92% of the edges in the younger mouse GCN are lost in the older mouse GCN, differences in network architecture may be attributed to age-related changes in key biological pathways in CD3+ splenocytes.

Another limitation of this study is that the dataset contains expression data from an array of T-cell subtypes not separated based on differentiation markers. Therefore, changes in co-expression behavior attributable to naïve, CD4+, or CD8+ cells cannot be discerned through this analysis.

In humans CD4, CD8, and CD3 T-cells present some overlap in gene expression during an immune response [73]. While thymic output of T-cells decreases with age, it is known that in some cases, older mice still produce naïve T-cells [25, 74]. This suggests the spleen dataset for the older mouse network may include expression data from both new and senescent T-cells, cell types which might demonstrate differential gene co-expression and proliferation propensity [30]. However, the production of naïve T-cells in mice is diminished around 5-10 fold in 18 old months mice, as it is also the case in human at midlife, at about 50 years [25]. In humans, there is 95% gene similarity between naïve and memory T-cells [75]. In C57BL/6 mice, both isolated naïve and memory CD4 T-cells from old mice present different alteration in gene expression, but both expression profiles are turned to inflammation [76]. Additionally, naïve and memory CD4 T-cells share fewer genes in common in old than in young mice, suggesting also a deterioration of these transcriptional pathways with aging. Over time, naïve T-cells may also increase their ability to persist [77]. Certain transcriptional changes, such as an increase in metabolism during the transition from naïve to effector T-cells, could not be accessed here [78].

The other limitation is the evolution of Treg cells, which represent only 1% of lymphocytes (about 5-10% of CD4), but their numbers increase with aging while their functionalities and regulatory functions decrease with the secretion of IL-2. In young mice, IL-2 at low doses stimulates Treg and negatively regulate T-cell proliferation, while at high doses Il-2 upregulates the proliferation of T-cells [25]. The role of Tregs in adipose-associated tissues is also correlated with increasing expression of anti-inflammatory genes such as IL-10.

Of course, these different age-related alterations in cell behavior are not specific to T-cell subtypes. Studies have found biomarkers of aging and senescence in other tissues in mice that suggest there may be cell-specific changes in gene expression with age [79]. Due to these variations, further investigations into co-expression in T-cells based on cellular age and T-cell subtype are necessary.

Interestingly, results by Vibert & Thomas-Vaslin show T-cell proliferation/death rates has higher inter-individual variation in the older mouse group when compared to the younger mouse group [24]. This may have an impact on the increased co-expression relationships evident in the old mouse GCN. Genes that are co-expressed in the old mouse group may also vary, resulting in the increased representation of nodes and edges in the old mouse GCN. This suggests co-expression pathways affected by age in CD3+ splenocytes vary based on genetic background and individual influences, demonstrating the need for future investigation into co-expression relationships in T-cells with age.

## Methods

### Mice

C57BL/6J (B6), BALB/cByJ and FVB/N (FVB) mice were obtained from Charles River Laboratories at 4 weeks of age, maintained in SPF conditions in our animal house (Centre d’Exploration Fonctionnelle Pitié Salpétrière–Paris) and fed with the same diet. Mice were sacrificed at 2 and 22-24 months of age. Mice were manipulated according to European council directive 86/609/EEC of 24 November 1986 and with the approval of an ethics committee.

### Sorting of CD3+ cells for mRNA preservation

Mice were sacrificed by cervical dislocation and spleens were collected. Cell suspensions were obtained by mechanical disruption of organs in PBS + 3% newborn calf serum at 4°C, were then washed, and viable cells were counted by trypan blue exclusion. CD3+ cells were isolated by positive selection using anti-CD3 biotine followed by streptavidin beads labelling and passage through LS magnetic Miltenyi column to recover fixed cells.

### Transcriptomics profiling

Gene expression was measured in CD3+ splenocytes extracted from FVB/N (n=4 young and n=3 old), C57BL/6N (n=4 young and n=3 old), and BALB/cByJ (n=4 young and n=5 old) mice. The young mice were age two months old. The old mice were age 22-24 months old.

One million purified cells were lysed in Trizol (Invitrogen) and immediately transferred to -80°C for storage. Samples were then lysed and total RNA was purified using Trizol (Invitrogen) according to the manufacturer’s instructions. RNA yield was assessed using a NanoDrop 1000 spectrophotometer (NanoDrop Products, Thermo Fisher Scientific). RNA integrity was assessed using an Agilent Bioanalyzer showing a quality RNA integrity number of 8–10 (Agilent Technologies). The RNA was processed using the Illumina TotalPrep RNA Amplification Kit Protocol according to the manufacturer’s protocol. Briefly, labeled complementary RNAs (cRNAs) were hybridized overnight with Illumina MouseWG-6 v2.0 Expression BeadChip arrays. The arrays were then washed, blocked, stained and scanned on an Illumina BeadStation following the manufacturer’s protocols. Illumina BeadStudio software (Illumina) was used to generate signal intensity values from the scans as previously described [85].

### Filtration and normalization of transcriptomics data

Microarray probes were filtered out from the analysis if their expression was below the detection limit (p-value < 0.05) in at least 2 out of 3 samples in both the microenvironment and control groups. Next, data were log-transformed and normalized by the quantiles method using the R package limma v3.28.4.

### Identification of specific molecular signatures

Specific molecular signatures were generated and statistically tested using the ICA/GSEA method [86], a strategy that combines the Independent Component Analysis (ICA) and the Gene Set Enrichment Analysis (GSEA). ICA, an unsupervised method, separates gene expression into non-Gaussian and statistically independent components. From each component, two potential molecular signatures were defined as those genes having extreme loading on components. Signatures were assembled in a database and tested for their enrichment in GSEA using the “pre-ranked list” option. Gene lists were sorted according to the value of the limma’s eBayes statistic. We used the weighted scoring scheme to compute the enrichment score. GSEA provides a normalized enrichment score, permutation p-value and FDR q-value indicating the significant level of each signature. A detailed explanation of GSEA can be found in Subramanian et al. [87]. Based upon altered gene expression in all aging mice, a total of 130 immune-related genes from the signature with the highest p-value (a signature named C2-5) was chosen for analysis among 900 signatures [24]. This signature was chosen since it allows to distinguish young from old mice, independently of their genetic origin. The resulting gene expression data was then grouped by age (12 young mice and 11 old mice) and gene co-expression.

### Gene co-expression networks construction

The Spearman rank correlations (*r*_s_) measuring co-expression levels were standardized by squaring *r*_s_, and the coefficient of determination R^2^ was used for network construction. This transformation ensures that the co-expression dataset included both strong positive and strong negative expression correlations. Significant co-expressed genes incorporated into network construction were defined as correlated gene pairs which had an R^2^ > 0.8 and a p-value < 0.01 [84]. The Python scripts used to filter this dataset based on these parameters are available on GitHub (http://github.com/mairml/network-analysis).

### Gene functional enrichment analysis

Human orthologues for the *mus musculus* genes, in the two networks, were determined using OrthoDB (http://orthodb.org). Additional functional annotations were retrieved from GenBank. All orthologues and functional annotations are provided in **S1 Table**.

MSigDB from GSEA was used to determine genes in each network that share common biological pathways in KEGG and REACTOME databases (http://software.broadinstitute.org/gsea/msigdb/annotate.jsp). Gene overlaps were calculated using an FDR of less than 0.05 in the Novartis human tissue compendium. Ingenuity Pathway Analysis was additionally utilized to analyze pathway enrichments in the dataset.

### Network topological analysis

Both the young and old mice networks were visualized using Cytoscape version 3.6.0 (http://cytoscape.org/download.php). The following network properties were calculated: network diameter, eigenvalue and betweenness centrality, clustering and closeness, network stress, network radius, network eccentricity, clustering, local influence, indirect and total influence. Built-in Cytoscape network analysis tools were used to find the mean shortest path, mean edge count, network diameter, radius and clustering coefficients for both GCNs. CentiScaPe version 2.2 was used to calculate centrality measures for each of the two networks (http://apps.cytoscape.org/apps/centiscape). The Grafman software was used to cross-check analysis results. The Grafman software is available from Bonchev [85]. Cytoscape plugin MCODE (http://apps.cytoscape.org/apps/mcode) was used to identify network clusters. Node influence was calculated using the algorithms discussed by Qiao *et al*. [60]. Adjacency matrices were calculated in Python (available at GitHub) and visualized in using the R package pheatmaps (http://cran.r-project.org/package=pheatmap). Distance matrices were calculated in Python (available at GitHub). Frequency distribution for each network was calculated using a Python script (available at GitHub). The frequency distributions were then scaled to probabilities and subsequently plotted in R v 3.1.1. Log-log plots associated with each of the model forms were used to estimate the model parameters. All parameters were estimated using linear regression on the log-transformed data [35–36].

## Acknowledgments

We would like to thank Dr. Phuong Pham (Sorbonne Université- INSERM UMRS 959 Immunology, Immunopathology, Immunotherapy CERVI, Pitié-Salpêtrière Hospital, Paris, France) for providing the C2-5 signature, for the conversion of microarray data into correlation matrices and for the constant support, and advice throughout the execution of this project. We would also like to thank Dr. Danail Bonchev (Virginia Commonwealth University) for his generous assistance in providing us with Grafman software as well as for his time in discussing network mathematics with us. Moreover, we wish to thank him for his assistance with some of the network calculations. Lastly, we would like to thank Dr. Wei Shan (Beihang University, China) and his team for his sharing their software for the calculation of local and global node power in a network.

## Supporting Information

**S1 Table. Summary of genes appearing in the age-related GCNs.** Genes with co-expression relationships represented in either or both the old and young networks along with their human ortholog and known functions.

**S2 Table. Network of 2-month old mouse CD3+ splenocytes.** All co-expression relationships appearing in the 2-month old mouse GCN are documented in this file.

**S3 Table. Network of 22/24-month old mouse CD3+ splenocytes.** All co-expression relationships appearing in the 22/24-month old mouse GCN are documented in this file.

**S1 Fig. Heatmap representation of gene expression signature from splenic T-cells obtained from young and old mice and across 3 mouse strains. The C2-5 signature with 130 selected genes was identified from ICA/GSEA.** The C2-5 signature distinguishes young from old mice (A), but also the genetic origins in young mice (B). The clustering of the 3 strains in young mice (B) is, however, lost in old mice (C). Key genes where co-expression was observed are underlined with TCR related genes (green boxes) co-stimulation molecules as CD28 (pink), CD69 an early activation marker and other markers revealed as centrality as the receptor for leptin (Leprotl1 gene).

**S2 Fig. Ingenuity Pathway Analysis of the C2-5 aging signature reveal altered gene expression and pathways in old mice.** (signature score: 40, focus molecules: 23). In A) genes with altered expression in the T-cells from old mice are involved in cellular development, cellular growth and proliferation, hematological system development and function. Genes belonging to TCR complex are underlined in green, CD28 downstream genes in orange, and those involved in T-cell signal transduction in blue. B) For this C2-5 aging signature, the top disease and functions are indicated.

**S3 Fig. MCODE generated network clusters.** In this figure we display only the MCODE generated clusters, not the full network structures. A total of four clusters were generated for the young group (A), and six clusters generated for the old group (B).

